# Centrioles generate two scaffolds with distinct biophysical properties to build mitotic centrosomes

**DOI:** 10.1101/2024.04.09.588708

**Authors:** Siu-Shing Wong, Joao M. Monteiro, Chia-Chun Chang, Min Peng, Nada Mohamad, Thomas L. Steinacker, Saroj Saurya, Alan Wainman, Jordan W. Raff

## Abstract

Mitotic centrosomes form when centrioles recruit large amounts of pericentriolar material (PCM) around themselves. The PCM comprises hundreds of proteins, yet it can assemble and disassemble within minutes, leading to much debate about its physical nature. Here we show that *Drosophila* Spd-2 fluxes out from centrioles, recruiting Polo and Aurora A to catalyse the assembly of a solid-like Cnn-scaffold and a more liquid-like TACC-scaffold, respectively. Both scaffolds can assemble and recruit PCM proteins independently, but both are required for proper centrosome assembly, with the Cnn-scaffold providing mechanical strength, and the TACC-scaffold locally concentrating centriole and centrosome proteins. Recruiting Spd-2, but not Cnn or TACC, to the surface of synthetic beads injected into early embryos reconstitutes several aspects of mitotic centrosome assembly on the bead surface, and this depends on the ability of Spd-2 to recruit Polo and AurA. Thus, Spd-2 molecules flux out from the centriole recruiting Polo and AurA to promote the assembly of two scaffolds with distinct properties to support mitotic centrosome assembly in flies.

## Introduction

Centrosomes are non-membrane bound organelles that form when centrioles recruit pericentriolar material around themselves (Bornens, 2021; Vasquez-Limeta and Loncarek, 2021; Gomes Pereira et al., 2021; Conduit et al., 2015). The PCM comprises several hundred proteins (Alves-Cruzeiro et al., 2014), and centrosomes function as important MT organising centres and cellular coordination hubs whose dysfunction has been linked to a plethora of human pathologies (Goundiam and Basto, 2021; Godinho and Pellman, 2014; Bettencourt-Dias et al., 2011; Nigg and Raff, 2009).

Despite its molecular complexity, the mitotic PCM can assemble and disassemble rapidly as cells prepare to enter and exit mitosis, respectively (Palazzo et al., 2000; Enos et al., 2018; Magescas et al., 2019; Mittasch et al., 2020)—prompting much debate about its biophysical nature (Rale et al., 2018; Woodruff, 2018; Raff, 2019; Lee et al., 2021; Woodruff, 2021). The mitotic PCM must be physically strong enough to resist the forces exerted by the spindle and astral MTs that it organises, but also provide an environment in which hundreds of proteins are concentrated and can potentially interact. In particular, there has been much debate about whether liquid-liquid phase separation (LLPS) may be important for mitotic centrosome assembly, as has been suggested for several other non-membrane bound organelles (Shin and Brangwynne, 2017; Alberti and Hyman, 2021).

In flies and worms a relatively simple pathway of mitotic PCM assembly has been proposed. The centriole and PCM protein Spd-2/SPD-2 (fly/worm nomenclature) (O’Connell et al., 2000; Kemp et al., 2004; Dix and Raff, 2007; Giansanti et al., 2008a) recruits Polo/PLK-1 (Decker et al., 2011; Alvarez Rodrigo et al., 2019; Alvarez-Rodrigo et al., 2021), which then phosphorylates Cnn/SPD-5 to stimulate the assembly of a macromolecular “scaffold” that recruits the many other PCM “client” proteins (Decker et al., 2011; Woodruff et al., 2015; Cabral et al., 2019; Ohta et al., 2021; Conduit et al., 2014a; b). Cnn and SPD-5 have no obvious sequence homology, but they are both large coiled-coil-rich proteins that can assemble into large scaffolding structures (Woodruff et al., 2017; Feng et al., 2017; Nakajo et al., 2022). This pathway appears to be widely conserved, and vertebrate homologues of Spd-2/SPD-2 (CEP192) (Zhu et al., 2008; Gomez-Ferreria et al., 2007; Joukov et al., 2010; Chinen et al., 2021), Polo/PLK-1 (PLK1) (Lane and Nigg, 1996; Haren et al., 2009; Lee and Rhee, 2011; Joukov et al., 2014; Meng et al., 2015) and Cnn (CDK5RAP2/CEP215) (Fong et al., 2008; Choi et al., 2010; Lizarraga et al., 2010; Barr et al., 2010; Tátrai and Gergely, 2022), have all been implicated in mitotic centrosome assembly.

The fly Cnn scaffold appears solid-like (Feng et al., 2017), but purified recombinant worm SPD-5 forms condensates *in vitro* that exhibit at least transient liquid-like properties, leading to the suggestion that the centrosome in worms is a condensate that forms through LLPS and concentrates MT organising proteins (Woodruff et al., 2017). Intriguingly, in mammalian oocyte spindles—which lack canonical centrioles and centrosomes—a different protein, TACC3, scaffolds a liquid-like spindle domain (LISD) that is essential for meiotic spindle assembly and that is also proposed to form via LLPS (So et al., 2019). The LISD is thought to be a specialised feature of meiotic spindles, as no LISD could be detected on mitotic spindles that were artificially induced to lack centrosomes (So et al., 2019). TACC proteins, however, are prominent components of mitotic centrosomes in many species (Gergely et al., 2000a; b) so we wondered whether centrosomes might organise a TACC-dependent LISD-like scaffold at normal mitotic spindles that form in the presence of centrosomes.

Here, we show that this is the case and that Spd-2 not only recruits Polo to stimulate the assembly of a solid-like Cnn scaffold, but also Aurora A (AurA) to stimulate the assembly of a more liquid-like TACC scaffold that appears to be analogous to the mouse oocyte LISD. The Polo/Cnn and AurA/TACC scaffolds recruit PCM client proteins independently, but both are required for efficient mitotic centrosome assembly in embryos—with the Cnn-scaffold providing mechanical strength, and the TACC-scaffold forming an extended structure around the centrioles that concentrates key centriole and centrosome proteins. We go on to show that centrioles generate an outward flux of Spd-2 molecules, and that recruiting Spd-2, but not Cnn or TACC, to the surface of synthetic beads injected into early embryos is sufficient to reconstitute several aspects of mitotic PCM assembly on the bead surface. Together, these studies demonstrate that Spd-2 acts as a nexus for centriole-driven mitotic centrosome assembly in fly embryos—fluxing outwards from the mother centriole to recruit Polo and AurA to stimulate the assembly of two independent scaffolds that together drive the assembly and outward expansion of the mitotic PCM. Importantly, although the TACC scaffold is more liquid-like than the Cnn-scaffold, it is not clear that the TACC scaffold is formed by LLPS.

## Results

### *Drosophila* TACC forms a centrosomal scaffold independently of the Cnn scaffold

To test whether centrosomes organise a TACC-dependent LISD-like scaffold we compared the centrosomal behaviour of GFP-TACC and GFP-Cnn in early *Drosophila* embryos. These embryos cycle rapidly between S-phase and mitosis with no Gap phases, so their centrioles essentially constantly organise a mitotic-like PCM (Wong et al., 2022). As reported previously, both GFP-TACC and GFP-Cnn spread out from the centrosome along the centrosomal MTs, often breaking off from the centrosome periphery as “flares” (Megraw et al., 2002; Lee et al., 2001) (*arrows*, Figure 1A, left panels). If MTs were depolymerised with colchicine, flaring was suppressed and GFP-Cnn formed a relatively inhomogeneous scaffold with irregular edges, while GFP-TACC formed a larger, more homogeneous, structure with smoother edges (Figure 1A, right panels). In colchicine-injected embryos the TACC structure extended significantly beyond the Cnn scaffold (Figure 1B), which could be observed directly in embryos co-expressing GFP-TACC and RFP-Cnn, (Figure 1C). This suggests that the TACC structure has some structural integrity independent of the Cnn scaffold, and we hereafter refer to it as a TACC scaffold.

**Figure 1.**
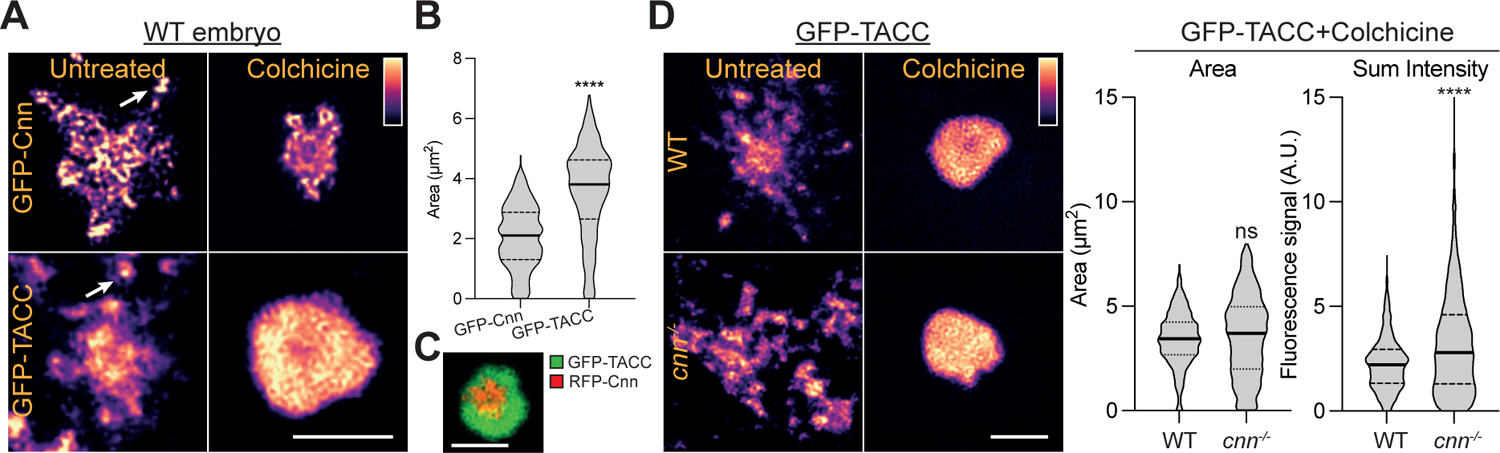
GFP-Cnn and GFP-TACC behave differently at centrosomes. (A) Images show the centrosomal distribution of GFP-Cnn or GFP-TACC in either untreated or colchicine-injected WT embryos. Arrows highlight “flares” breaking away from the main centrosome scaffold. (B) Violin plot shows the area (Median±Quartiles) of the GFP-Cnn and GFP-TACC scaffolds in colchine-injected WT embryos. N=13-20 embryos; n=∼400-800 centrosomes. (C) Image shows the distribution of RFP-Cnn and GFP-TACC at a typical centrosome in a colchicine-injected WT embryo. (D) Images show the distribution of GFP-TACC in either untreated or colchicine-injected WT or *cnn^-/-^* embryos. Violin plots show the area or fluorescence-intensity (Median±Quartiles) of the centrosomal GFP-TACC in these embryos N=10-16 embryos; n=∼700-1300 centrosomes for each genotype. Scale bars = 2μm. Statistical comparisons used Mann-Whitney test (****: P<0.0001; ns: not significant).

To test whether the TACC scaffold assembled upon the Cnn scaffold, we expressed GFP-TACC in embryos laid by c*nn*^-/-^ mutant females (hereafter *cnn^-/-^* embryos). GFP-TACC was still detected at centrosomes in *cnn^-/-^*embryos, but it formed more extensive flares that spread out over a broader area (Figure 1D, left panels; Video 1). When flaring was suppressed by colchicine injection, however, the recruitment of GFP-TACC to centrosomes was, if anything, slightly enhanced in *cnn^-/-^*embryos (Figure 1D). Moreover, a fluorescence recovery after photobleaching (FRAP) analysis revealed that GFP-TACC was recruited to centrosomes with similar kinetics in both WT and *cnn^-/-^* embryos (Figure S1). Thus, the TACC scaffold can form independently of the Cnn scaffold, but it is normally stabilised by the Cnn scaffold, which helps prevent its dispersion along the centrosomal MTs.

### The Cnn and TACC scaffolds have different biophysical properties

We next used FRAP to compare the behaviour of the Cnn- and TACC-scaffolds. In colchicine-injected embryos, centrosomal GFP-Cnn fluorescence appeared to recover slowly and preferentially around the central region of the centrosome, while GFP-TACC fluorescence recovered more quickly and more evenly throughout the centrosomal region (Figure 2A; Video 2). The duplicated centrosomes often only partially separated in these colchicine-injected embryos. In this situation, GFP-Cnn usually formed discrete scaffolds around each centrosome, with a clear boundary between the two centrosomes that was stable over time (arrows, Figure 2B); in contrast, GFP-TACC usually formed a more continuous scaffold with no clear demarcation between the two centrosomes (Figure 2B; Video 3). Kymographs of partially photobleached centrosome pairs revealed that GFP-Cnn molecules in the scaffold appeared immobile, and fluorescence recovered at the bleached centrosome independently of the non-bleached centrosome; in contrast, GFP-TACC molecules were mobile, and readily diffused from the non-bleached centrosome to the bleached centrosome (Figure 2C; Video 4). Moreover, in *cnn^-/-^* embryos injected with colchicine we often observed GFP-TACC flares “fusing” back with the main centrosomal GFP-TACC scaffold, a behaviour we did not observe with similarly positioned GFP-Cnn flares (Figure 2D). Thus, the Cnn scaffold exhibits more solid-like, and the TACC scaffold more liquid-like, behaviour.

**Figure 2.**
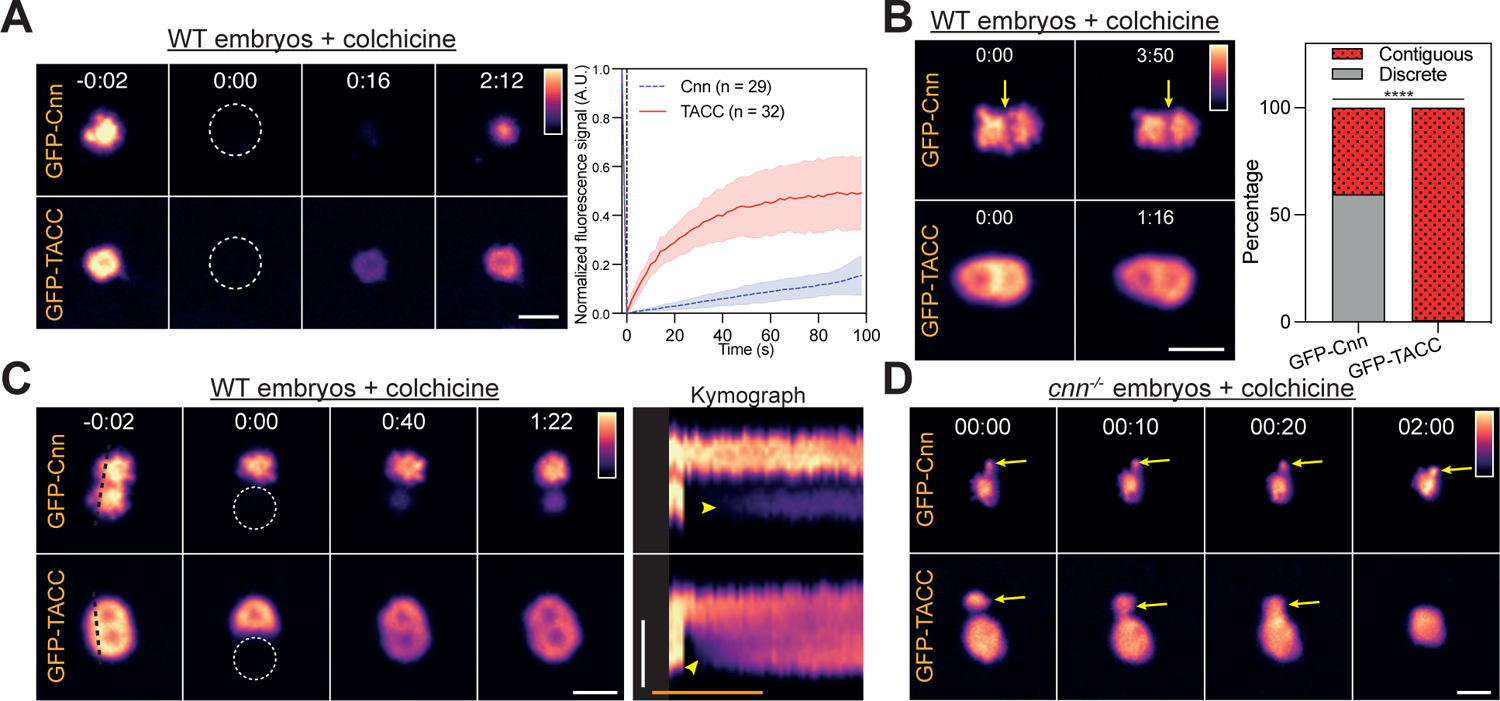
Dynamic analysis of the GFP-Cnn and GFP-TACC scaffolds. (A) Images show the behaviour of GFP-Cnn or GFP-TACC in a FRAP experiment in WT colchicine-injected embryos. Time (mins:secs) is indicated; centrosomes were bleached at t=0:00. Graph shows each protein’s normalised fluorescence intensity recovery profile (Mean±SD). N=6-9 embryos; n=∼20-40 centrosomes. (B) Images show the distribution of GFP-Cnn or GFP-TACC at two closely abutted centrosomes that failed to separate properly in colchicine-injected WT embryos. GFP-Cnn usually occupies discrete centrosomal domains separated by a clear demarcation (*arrow*) that is stable over time. GFP-TACC usually forms a single continuous domain surrounding both centrosomes. Bar chart quantifies the percentage of centrosomes with discreet or continuous domains. N=12-20 embryos; n=100-300 centrosomes. Contingency significance (i.e. the significance of the difference in the proportion between the two groups) was calculated using a Fisher’s exact test (****: P<0.0001) (C) Images show the FRAP behaviour of GFP-Cnn or GFP-TACC at closely-paired centrosomes in which only one centrosome has been bleached (at t=0:00). The Kymograph shows how fluorescence intensity changes over time along the black-dotted lines indicated in the pre-bleach images (t=-0:02): white scale bars = 2μm; cyan scale bar = 2min. (D) Images show how flares (*arrows*) of GFP-Cnn do not detectably fuse with the main centrosome scaffold, but such fusion events are readily detected with flares of GFP-TACC.

### The centrosomal TACC scaffold is related to the mouse meiotic spindle LISD

In the mouse meiotic spindle, TACC3, AurA and Clathrin Heavy Chain (CHC) are required for LISD assembly (So et al., 2019; Wang et al., 2021; Blengini et al., 2021). To test whether these constituents were required for the assembly of the fly centrosomal TACC scaffold we quantified GFP-TACC scaffold assembly in embryos laid by mothers in which we halved the genetic dosage of either *Tacc*, *aurA* or *Chc*. These experiments were performed in *cnn^-/-^* embryos injected with colchicine to avoid any potentially confounding effects that halving the dose of these proteins might have on the Cnn scaffold, and vice versa. All these perturbations significantly reduced the size of the TACC scaffold (Figure 3A). In contrast, halving the genetic dose of the Pericentrin-like protein (Plp) or Plk4—proteins that are not components of the mouse oocyte LISD (So et al., 2019) but have a prominent role in centrosome (Richens et al., 2015; Lerit et al., 2015) and centriole (Bettencourt-Dias et al., 2005; Aydogan et al., 2020) assembly, respectively—had no effect on TACC scaffold size (Figure 3). Thus, the centrosomal TACC scaffold in flies appears to be closely related to the TACC3-dependent LISD in mouse oocytes.

**Figure 3.**
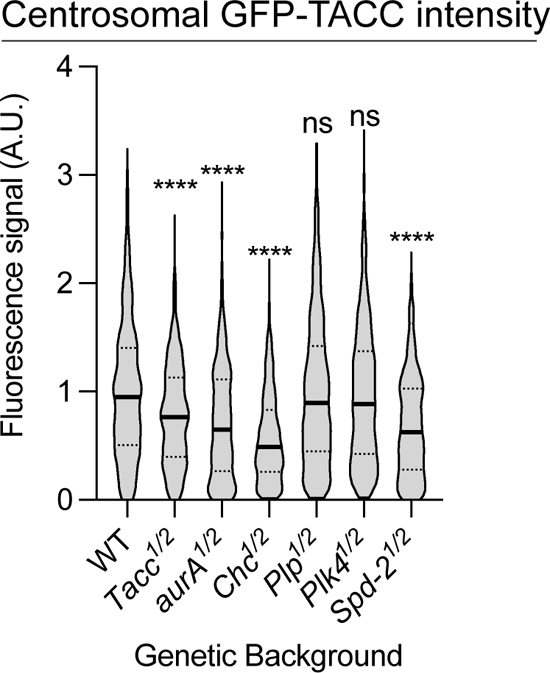
Analysis of the molecular requirements for TACC-scaffold assembly. Violin plot quantifies the centrosomal fluorescence intensity (Median±Quartiles) of centrosomal GFP-TACC in living embryos laid by mothers that are heterozygous for null mutations in the indicated genes (so the embryos contain approximately half the normal amount of each protein—see Materials and Methods). N=10-15 embryos; n=500-1000 centrosomes. Statistical significance was calculated using a Kruskal-Wallis test, followed by a Dunn’s Multiple Comparison (****: P<0.0001; ns: not significant).

### AurA phosphorylates TACC to promote TACC scaffold assembly

AurA/AURKA phosphorylates TACC/TACC3 on a conserved Serine, S863/S558 in flies/humans (Barros et al., 2005; Kinoshita et al., 2005), which allows TACC3 to interact with CHC (Lin et al., 2010; Booth et al., 2011; Hood et al., 2013). We reasoned, therefore, that AurA could promote TACC scaffold assembly by phosphorylating TACC-S863 to stimulate the TACC/CHC interaction. To test this possibility, we expressed either WT GFP-TACC or a non-phosphorylatable GFP-TACC-S863L mutant (Barros et al., 2005) in embryos co-expressing RFP-Cnn. GFP-TACC-S863L was expressed at slightly higher levels than WT GFP-TACC (Figure S2A) but, although the mutant protein was still recruited to centrosomes, it did not detectably form an independent scaffold that extended beyond the Cnn scaffold, so the total amount of centrosomal TACC-S863L was dramatically reduced (Figure 4A). Moreover, a FRAP analysis of mNeonGreen (NG)-TACC-WT and NG-TACC-S863L expressed in embryos laid by *Tacc^-/-^* mutant females (*Tacc^-/-^*embryos) (Figure S2B), revealed that the mutant protein turned over much faster at centrosomes than the WT protein, consistent with the hypothesis that TACC-S863L cannot efficiently form a scaffold (Figure 4B). We conclude that AurA phosphorylates TACC-S863 to promote TACC-scaffold assembly, presumably by promoting the TACC/CHC interaction. Interestingly, Cnn-scaffold size appeared largely unperturbed in embryos expressing GFP-TACC-S863L, suggesting that the Cnn scaffold can assemble independently of the TACC scaffold (Figure 4A).

**Figure 4.**
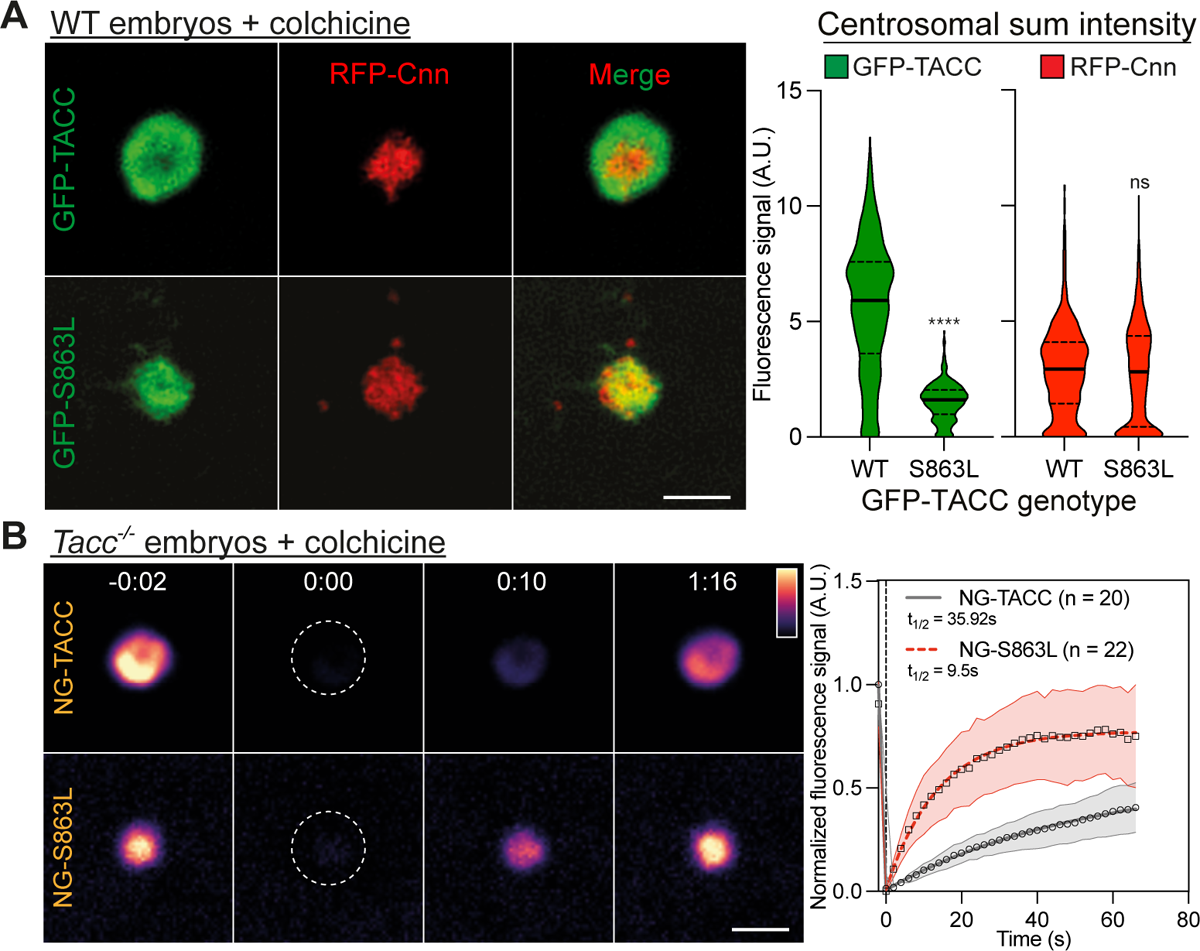
TACC phosphorylation on Ser863 promotes scaffold assembly. (A) Images show the centrosomal localisation of WT GFP-TACC or the GFP-TACC-S863L mutant in WT embryos expressing RFP-Cnn. Violin plots quantify the total centrosomal fluorescence intensities (Median±Quartiles) of GFP-TACC and GFP-S863L (left) or RFP-Cnn (right) in these embryos. N=10 embryos; n=800-900 centrosomes for each group. Statistical comparisons used a Mann-Whitney test (****: P<0.0001; ns: not significant). (B) Images show the behaviours of NG-TACC or NG-S863L in a FRAP experiment in colchicine-injected *Tacc^-/-^* embryos. Time (mins:secs) is indicated; centrosomes were bleached at t=0:00. Graph shows each protein’s normalised fluorescence intensity recovery profile (Mean±SD). Fitted lines and their parameters were generated using a One-Phase Association Model (Graphad Prism). N=4-9 embryos; n=20-30 centrosomes for each group. Scale bars = 2μm.

### Spd-2 helps recruit AurA and Polo to centrosomes

It has previously been shown that CEP192 proteins in humans and frogs help to recruit both PLK1 and AURKA to centrosomes (Joukov et al., 2014; Meng et al., 2015), so we wondered whether *Drosophila* Spd-2 might also recruit AurA to promote TACC scaffold assembly. In support of this possibility, halving the genetic dose of *Spd-2* reduced TACC scaffold assembly in a similar manner to halving the dose of *AurA* (Figure 3). In vertebrates, the interaction between CEP192 and AURKA is well characterised (Joukov et al., 2010, 2014; Meng et al., 2015) and the crystal structure of an interaction interface has recently been described (Park et al., 2023). It is unclear, however, if invertebrate versions of these proteins interact (Joukov and De Nicolo, 2018). We therefore used ColabFold-adapted AlphaFold2 to screen for potential interactions between *Drosophila* Spd-2 and AurA and found a reasonably high-confidence prediction (iPTM=0.59) between Spd-2229-310 and the AurA kinase domain (AurA155-421) (Figure 5A; Figure S3). This interaction comprised two independent interfaces involving two short regions of Spd-2 (Spd-2229-250 and Spd-2291-310), which we term **AurA b**inding **d**omain (ABD) 1 and 2, respectively, wrapping around the surface of the kinase domain. Interestingly, Spd-2-ABD1 and Spd-2-ABD2 are predicted to bind to similar regions on AurA/AURKA as human CEP192 (Park et al., 2023) and human TPX2 (Bayliss et al., 2003), respectively (Figure 5A; Figure S3).

**Figure 5.**
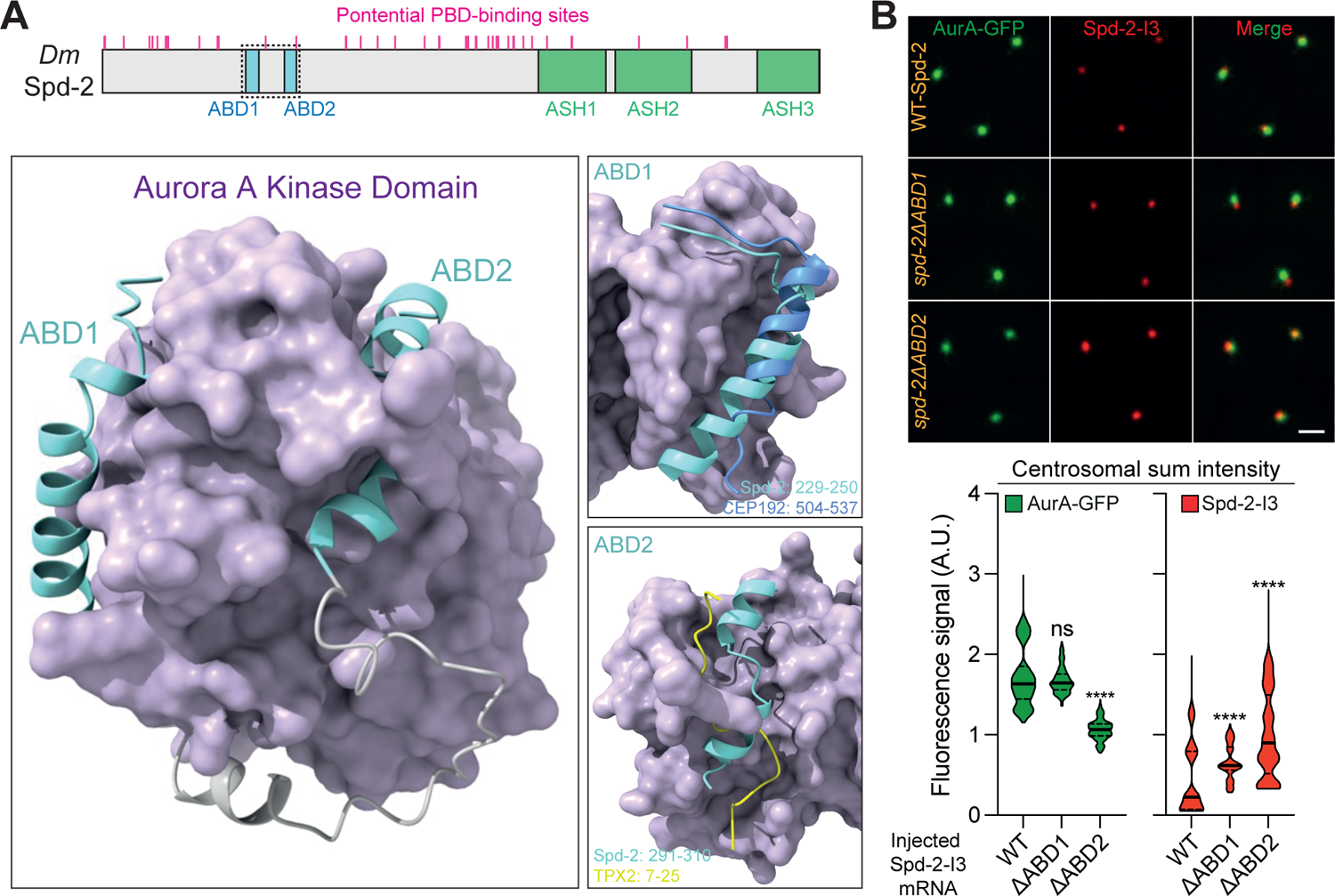
Identifying and testing potential interactions between Spd-2 and AurA. (A) Schematic representation of Drosophila Spd-2 highlighting the position of the two potential AurA binding domains (ABD1 and ABD2) (*cyan boxes*), the 3 C-terminal ASH domains (*green boxes*), and the multiple potential Polo-Box binding motifs examined previously (Alvarez Rodrigo et al., 2019) (*magenta lines*). The fragment of Spd-2 shown in the predicted structure below is indicated by a black dotted box. (B) ColabFold predicted structure of Spd-2_229-310_ (ABD1 and ABD2 in *cyan* with *grey* linker region) bound to the AurA kinase domain (*indigo* surface). The two putative interaction sites (ABD1_229-250_ and ABD2_291-310_) are shown in the boxes on the right overlayed with the previously described crystal structure of either human CEP192_504-537_ (*blue*, top box) or Xenopus TPX-27-25 (*yellow*, bottom box) bound to AURKA (overlayed here on the Drosophila AurA structure). See Figure S3 for a more detailed analysis. (C) Images show, and violin plots quantify (Median±Quartile), centrosome fluorescence intensity in embryos expressing AurA-GFP and injected with mRNA encoding mScarlet-I3-fusions to either WT-Spd-2, or mutant forms of Spd-2 in which the putative ABD’s have been deleted (Spd-2ΔABD1 and Spd-2ΔABD2). N= 9 embryos; n = 400-500 centrosomes for each group. Statistical significance was calculated using a Kruskal-Wallis test, followed by a Dunn’s Multiple Comparison (****: P<0.0001; ns: not significant).

To test whether ABD1 or ABD2 normally help to recruit AurA to centrosomes in flies we injected mRNA encoding mScarlet-I3-(I3)-fusions (Gadella et al., 2023) to either WT Spd-2 or forms of Spd-2 in which either ABD1 or ABD2 were deleted into embryos expressing AurA-GFP. In this assay, the injected mRNA is quickly translated, and the expressed I3-fusions compete with the endogenous unlabelled protein to bind to the centrosome (Novak et al., 2016). The mutant Spd-2-I3-fusions were recruited to centrosomes at higher levels than the WT Spd-2-fusions (for unknown reasons), but the centrosomal accumulation of AurA-GFP was either largely not affected (Spd-2ΔABD1) or was significantly reduced (Spd-2ΔABD-2) (Figure 5B). Thus, both the ABD1 and ABD2 deletions reduce the amount of AurA recruited to centrosomes (per molecule of Spd-2) with ABD2 appearing to have the more dominant role.

We have previously shown that Polo binds to Spd-2 via the Polo-Box domain (PBD) (Alvarez Rodrigo et al., 2019), which binds to phosphorylated S-S/T(P) motifs (Elia et al., 2003). A form of Spd-2 in which all of the potential PBD-binding S-S/T motifs (indicate by *magenta* lines in Figure 5A) have been mutated to T-S/T (previously referred to as Spd-2-ALL, but here called Spd-2ΔPolo, for simplicity), cannot bind the PBD and appears unable to recruit Polo to centrosomes (Alvarez Rodrigo et al., 2019). We confirmed that a Spd-2ΔPolo-I3 fusion also strongly reduced Polo-recruitment compared to WT Spd-2-I3 in the mRNA injection assay described above (Figure S4). Thus, as in vertebrates, *Drosophila* Spd-2 helps to recruit both Polo and AurA to centrosomes.

### Spd-2 molecules incorporate into the PCM close to the centrioles and then flux outwards

In colchicine-injected *cnn^-/-^* embryos co-expressing Spd-2-RFP and AurA-GFP, both proteins were concentrated around the centrioles but also spread throughout the TACC-scaffold region (Figure 6A, t= −0:02). A FRAP analysis, however, revealed that Spd-2-RFP fluorescence first recovered at the centrioles and then seemed to spread outwards through the TACC scaffold, while AurA-GFP fluorescence recovered more quickly both around the centriole and throughout the TACC scaffold (Figure 6A,B). This suggests that Spd-2 molecules initially bind close to the centriole and then flux outwards, recruiting AurA to initiate the assembly of the TACC scaffold—just as we have previously hypothesised that Spd-2 fluxes out from the centriole to recruit Polo to initiate the assembly of the Cnn scaffold (Conduit et al., 2014b, 2015). In support of this possibility, we observed several instances in colchicine-injected *cnn^-/-^* embryos where the TACC scaffolds became separated from the Spd-2-RFP-generating centrioles; this led to the generation of new TACC-scaffold around the centrioles, and to the eventual disassembly of the TACC-scaffolds that had lost their connection to the centrioles (Figure S5).

**Figure 6.**
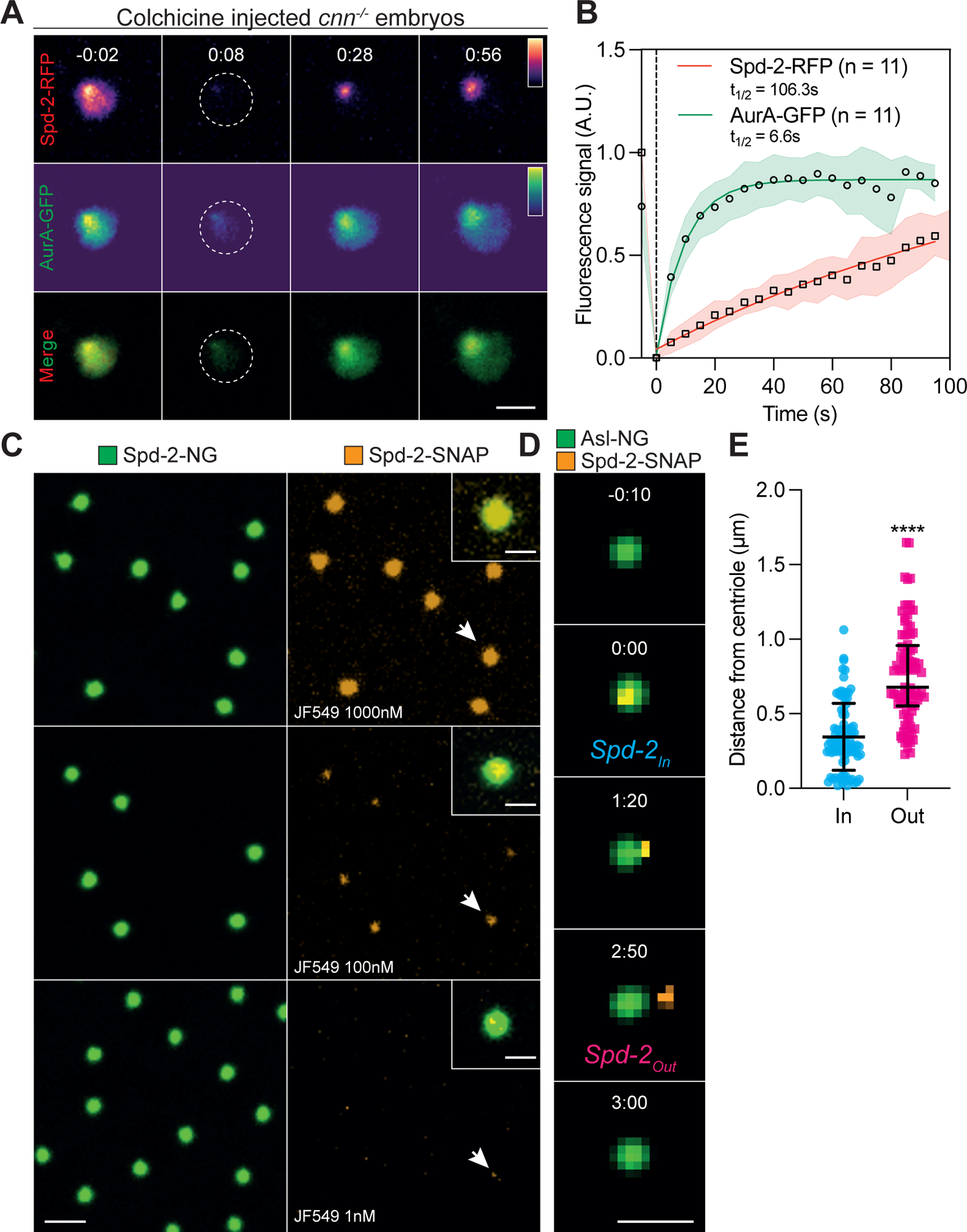
Spd-2 molecules flux outwards from the centrioles. (A) Images show the behaviour of Spd-2-RFP and AurA-GFP in a FRAP experiment in colchicine-injected *cnn^-/-^* embryos. Time (mins:secs) is indicated; centrosomes were bleached at t=0:00 (not shown). (B) Graph shows each protein’s normalised fluorescence intensity recovery profile (Mean±SD). N=9 embryos; n=11centrosomes. (C) Images show the centrosomes in embryos co-expressing Spd-2-mNG and Spd-2-SNAP and injected with different concentrations of the JF-549 SNAP-ligand. The lower the concentration of injected-ligand, the sparser the labelling of the centrosomes. The centrosomes highlighted with white arrows are shown in the enlarged insets. (D) Images show stills from a movie tracking a single molecule of fluorescently labelled Spd-2-SNAP binding to, and then unbinding from, a mother centriole (labelled with Asl-mNG). Time (min:secs) relative to the first detection of the single particle at the centriole (t=0:00) is indicated. (E) Scatter plot shows the distance (Median±Quartiles) between single molecules of fluorescently labelled Spd-2-SNAP and the centre of the mother centriole at the time the molecules initially bind to the centrosome (Spd-2In) and at the time they leave the centrosome (Spd-2Out) N=89 single molecules. Scale bars: (A,D) = 2μm; (C) = 4μm (inset = 2μm).

The outward flux of Spd-2, however, remains unproven, and the pattern of Spd-2 fluorescence recovery could be explained by the differential turnover-rate of Spd-2 binding sites spread throughout the PCM, rather than by an outward flux of Spd-2 molecules. Moreover, no outward SPD-2 flux could be detected at centrioles in *C. elegans* embryos (Cabral et al., 2019). To directly test our Spd-2-flux hypothesis in flies, we examined the behaviour of individual fluorescently-tagged Spd-2 molecules in living embryos. Such studies have provided great insight into the behaviour of proteins in their native environment (Nguyen et al., 2023), so we generated transgenic lines expressing Spd-2 fused to a SNAP-tag—a protein that is not fluorescent but which can be covalently coupled to a fluorescent SNAP-ligand (Keppler et al., 2003). We injected the yellow-fluorescent SNAP ligand JF-549 (Grimm et al., 2015) into *Drosophila* embryos expressing Spd-2-SNAP and Spd-2-NG. At an injection concentration of 1000nM, the SNAP-ligand appeared to label essentially all of the Spd-2-SNAP molecules, but the fraction of molecules labelled decreased as the injection concentration of the ligand decreased (Figure 6C). At a concentration of 1nM, only a small fraction of the centrosomes (usually 5-10%) were labelled, suggesting that any yellow-fluorescent signal at a centrosome was likely to be generated by a single fluorescent Spd-2-SNAP molecule (arrow, bottom panel, Figure 6C).

We then injected 1nM JF-549 into embryos laid by mothers expressing Spd-2-SNAP and Asl-NG—a mother centriole marker (Novak et al., 2014)—and imaged the embryos at 10sec intervals. We computationally identified single molecule-binding events (see Materials and Methods) (Figure 6D), and measured the distance between the mother centriole and each Spd-2-SNAP molecule at each timepoint that the molecule was bound to the centrosome (Figure 6E,F). On average, Spd-2-SNAP molecules bound into the PCM (*Spd-2*In) close to the centriole (345±225nm) (Mean±SD), moved outwards, and eventually unbound from the PCM (*Spd-2*Out) further away from the centriole (762±322nm) (Video 5). Thus, Spd-2 molecules exhibit an overall outward flux from the centriole in fly embryos.

### The TACC scaffold concentrates centrosome proteins

In mouse oocytes, the LISD helps to recruit proteins to the poles of the meiotic spindle (So et al., 2019). To test if the fly TACC scaffold functioned similarly, we compared the centrosomal enrichment of several fluorescent-fusions to centrosomal proteins in WT and *cnn^-/-^* embryos injected with colchicine (Figure 7A). All the proteins we tested were significantly enriched at centrosomes in WT embryos, and also within the TACC scaffold in *cnn^-/-^* embryos. Interestingly, Spd-2-NG, AurA-GFP and γ-tubulin GFP were more enriched at centrosomes in WT embryos than within the TACC scaffold in *cnn^-/-^* embryos, suggesting that the Cnn scaffold normally helps to recruit these proteins to centrosomes. In contrast, Msps-GFP, Klp10A-GFP and CHC-GFP were approximately equally enriched at centrosomes in WT embryos and in the TACC scaffold in *cnn^-/-^*embryos, indicating that the TACC scaffold may play the dominant part in recruiting these proteins to centrosomes even when the Cnn scaffold is present. This is perhaps to be expected for CHC, which appears to be a crucial component of the TACC scaffold and the LISD. We also examined the recruitment of two mutant forms of Cnn that do not support efficient Cnn-scaffold assembly (GFP-CnnΔLZ and GFP-CnnΔCM2) (Feng et al., 2017). Both were enriched in the TACC scaffold to a similar extent in colchicine-injected WT and *cnn^-/-^* embryos (Figure 7B), indicating that the TACC scaffold can concentrate Cnn molecules at centrosomes independently of the Cnn scaffold, and independently of the ability of the Cnn molecules to form a Cnn-scaffold.

**Figure 7.**
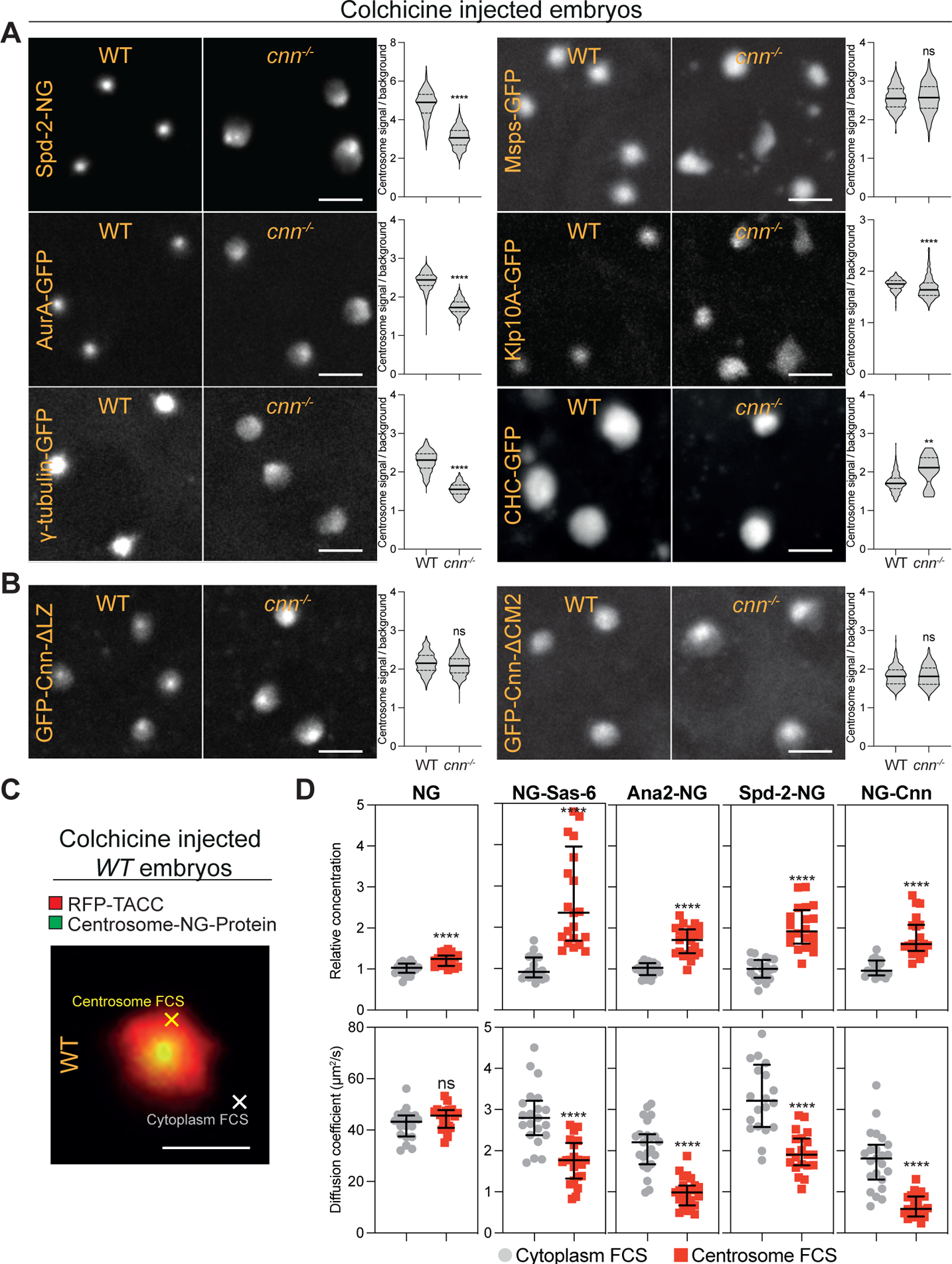
The TACC scaffold concentrates centriole and centrosome components. (A) Images show the accumulation of fluorescent-fusions to various proteins at centrosomes in WT or *cnn^-/-^* embryos treated with colchicine. Violin plots quantify (Median±Quartiles) the fold enrichment of the proteins at the centrosome compared to the cytoplasm. N=3-10 embryos; n=60-1000 centrosomes for each group. (B) These panels show the same as in (A) but for GFP-fusions to Cnn deletion mutants (ΔLZ or ΔCM2) that cannot form a Cnn scaffold. N=10-11 embryos; n=400-900 centrosomes for each group. (A,B) (C) Image shows a centrosome in an embryo expressing RFP-TACC and Ana2-NG and injected with colchicine. This illustrates the typical areas that were analysed by FCS in the outer regions of the centrosome (*yellow cross*) or the nearby cytoplasm (*white cross*). Scale bar = 2μm. (D) Scatter plots show the FCS-measured concentration (top plots) or diffusion rate (bottom plots) (Median±Quartiles) of NG or various NG-fusions in the cytoplasm (*grey circles*) or in the outer regions of the centrosome (*red squares*) in WT embryos injected with colchicine. N=20-25 embryos. Statistical significance was calculated using a Mann-Whitney test (****: P<0.0001; ns: not significant).

We next used Fluorescence Correlation Spectroscopy (FCS) to compare the concentration and diffusion rate of the centrosome building blocks Spd-2-NG and NG-Cnn within the cytoplasm (*white cross*, Figure 7C) and within the outer TACC-scaffold region that extends beyond the Cnn-scaffold (*yellow cross*, Figure 7C). As the PCM can also promote centriole assembly (Dammermann et al., 2004), we also examined the behaviour of the centriole building blocks Sas-6-NG and Ana2-NG (Arquint and Nigg, 2016; Gönczy and Hatzopoulos, 2019). We performed this analysis in colchicine-injected embryos to prevent any flares of centrosomal proteins moving into the cytoplasmic regions during the analysis period. As a control, we first analysed unfused-NG molecules; these were slightly enriched in the TACC-scaffold region, but their diffusion rate within the scaffold was not significantly altered compared to the cytoplasm (Figure 7D). In contrast, all the centriole and centrosome proteins were enriched in the TACC scaffold to a greater extent than NG, and their diffusion rates within the TACC-scaffold were significantly reduced (Figure 7D). Importantly, we obtained very similar results when we analysed the behaviour of some of these protein in *cnn^-/-^* embryos, indicating that in these experiments we were likely to be assessing the behaviour of proteins in the TACC scaffold, independently of the Cnn scaffold (Figure S6). We conclude that the TACC scaffold can concentrate key centriole and centrosome proteins, at least in part, by slowing their diffusion within the TACC scaffold region.

### Reconstituting mitotic PCM assembly on the surface of Spd-2-coated synthetic beads

Our data so far suggests a simple scheme whereby Spd-2 fluxes out from centrioles recruiting Polo and AurA to allow them to phosphorylate Cnn and TACC, respectively, to initiate scaffold (and so ultimately mitotic-PCM) assembly (Figure 8A). To test this scheme, we individually tethered Spd-2-GFP, GFP-Cnn or GFP-TACC to synthetic beads in embryos to see whether simply concentrating these proteins on the bead-surface was sufficient to initiate scaffold assembly (Figure 8B). We injected streptavidin-coated beads bound to a biotinylated GFP-Nanobody together with mRNA encoding either GFP, GFP-Cnn, GFP-TACC, or Spd-2-GFP into embryos expressing either RFP-Cnn or mCherry-TACC (to monitor scaffold assembly) or Jupiter-mCherry (to monitor MT organisation).

**Figure 8.**
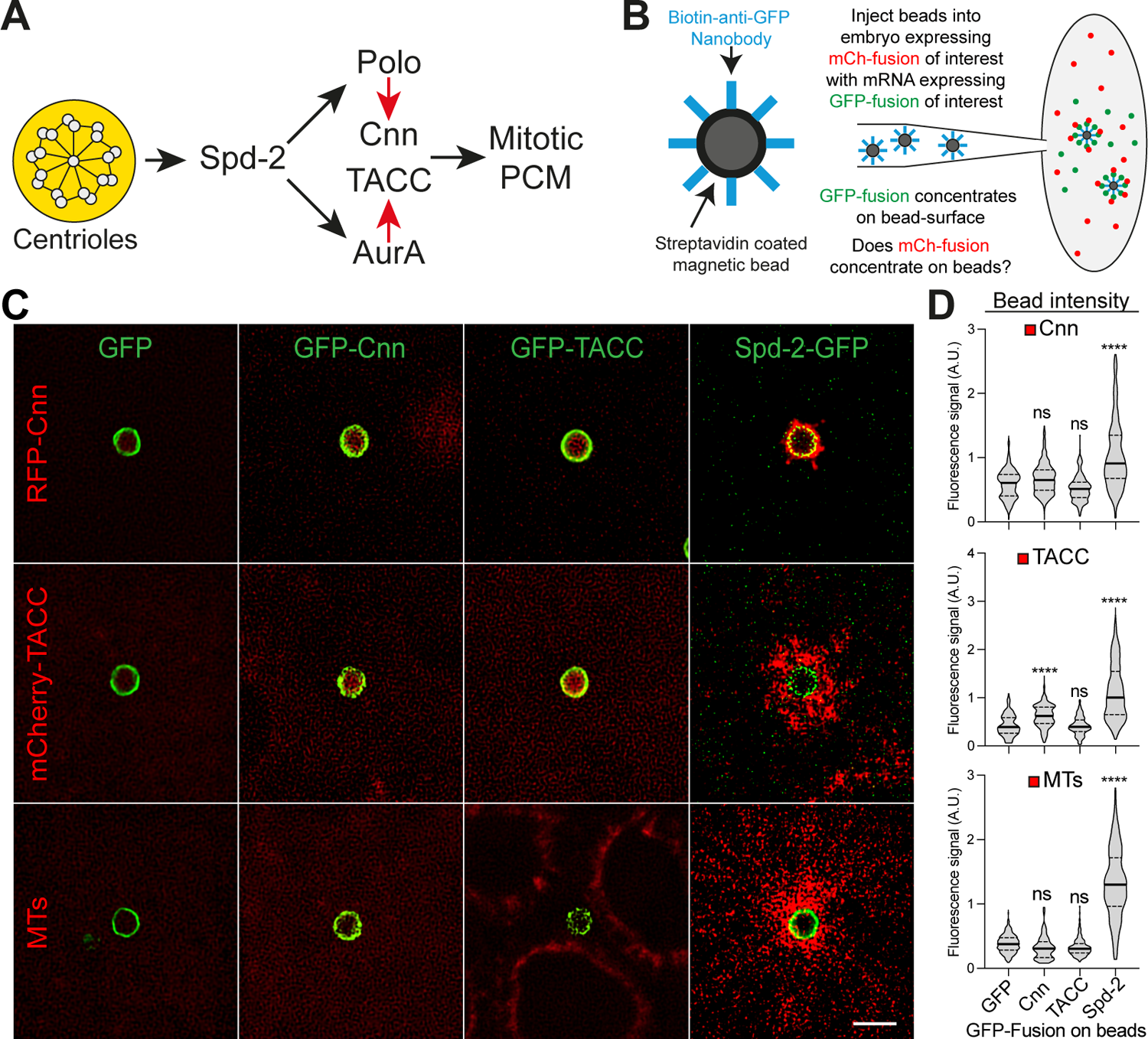
Reconstituting aspects of mitotic PCM assembly on the surface of synthetic beads. (A) Schematic illustrates the proposed pathway of mitotic-PCM-scaffold assembly. Centrioles provide a source of Spd-2 that fluxes outwards, recruiting AurA and Polo, which then phosphorylate TACC and Cnn (red arrows), respectively, to initiate the assembly of scaffolds that support the assembly of the mitotic PCM. (B) Schematic illustrates the synthetic bead injection assay. (C, D) Images show, and violin plots quantify (Median±Quartiles), the red-fluorescent signal—either RFP-Cnn, RFP-TACC (as markers of the PCM-scaffolds) or Jup-mCherry (as a marker of MTs)—on the beads when the beads are bound to GFP or the various GFP-fusions. Statistical significance was calculated by an Ordinary one-way ANOVA, followed by a Tukey’s Multiple Comparison (****: P<0.0001; ns: not significant). Note that, for unknown reasons, the GFP-Cnn bound beads appear to recruit significantly more RFP-TACC than controls, but this recruitment is not nearly as strong as that observed with the Spd-2-GFP coated beads.

In these experiments, the beads invariably recruited the GFP or GFP-fusion of interest, but only the beads bound to Spd-2-GFP appreciably recruited RFP-Cnn, mCherry-TACC and MTs (Figure 8C,D). The dynamics of the MTs organised by these beads cycled in synchrony with the dynamics of the MTs organised by the endogenous centrosomes, suggesting that the PCM scaffold assembled on the Spd-2-GFP bead surface can respond to cell-cycle cues (Video 6). Interestingly, the TACC scaffold significantly extended outwards around the bead, but the Cnn scaffold remained tightly associated with the bead-surface (Figure 8C,D). This suggests that the outward-flux of Spd-2 molecules from the centrioles, which presumably does not occur on the Spd-2-GFP-coated beads (as the Spd-2-GFP is bound directly to the high-affinity anti-GFP-nanobody), may be essential for the outward expansion of the solid-like Cnn-scaffold, but not for the expansion of the liquid-like TACC-scaffold.

To test whether the recruitment of Polo and/or AurA was required to allow the Spd-2-coated beads to organise the Cnn and/or TACC scaffolds, we adapted the beads assay to use streptavidin-coated beads bound to a biotinylated anti-ALFA-tag-Nanobody. This Nanobody binds to the 13aa ALFA-tag peptide (Götzke et al., 2019), and we co-injected mRNA encoding WT or various mutant ALFA-tagged versions of Spd-2 into embryos expressing either Polo-GFP and RFP-Cnn, or AurA-GFP and mCherry-TACC. Beads bound to WT Spd-2-ALFA strongly recruited all of these proteins, but beads bound to a form of Spd-2 that cannot recruit Polo—Spd-2-ΔPolo; previously referred to as Spd-2-ALL (Alvarez Rodrigo et al., 2019)—recruited essentially no Polo-GFP or RFP-Cnn, but recruited normal, or even slightly enhanced, amounts of AurA-GFP and mCherry-TACC (Figure 9). Conversely, when bound to Spd-2-ALFA mutants that perturb the interaction with AurA (Spd-2-ΔABD1 or Spd-2-ΔABD2), the ability of the beads to recruit AurA and TACC was severely diminished (Figure 9). Interestingly, the Spd-2-ΔABD1 coated beads recruited near-normal levels of Polo-GFP and RFP-Cnn, consistent with the idea that the two scaffolds are largely independent on one another, but the Spd-2-ΔABD2 coated beads recruited significantly reduced levels or Polo-GFP and RFP-Cnn (Figure 9). The reason for this difference between Spd-2-ΔABD1 and Spd-2-ΔABD2 is unclear, but we suspect that Spd-2-ABD2 may bind to and activate AurA in a way that allows AurA to help activate Polo.

**Figure 9.**
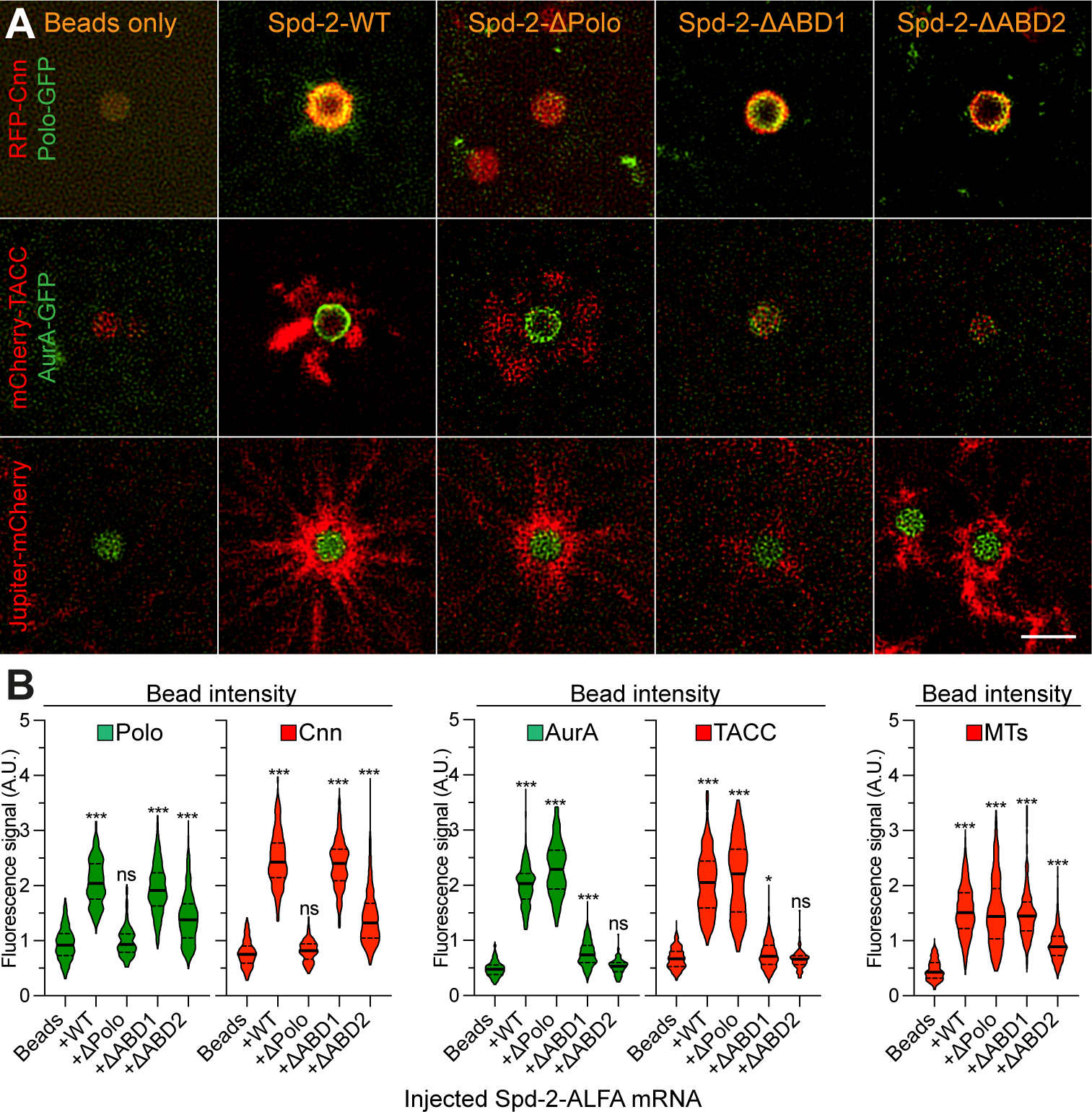
Spd-2 can recruit Polo/Cnn and AurA/TACC at least partially independently to organise PCM and MTs at synthetic beads. (A) Images show ALFA-tag-Nanobody-coupled synthetic beads that have been injected into embryos expressing either RFP-Cnn and Polo-GFP (top), mCherry-TACC and AurA-GFP (middle), or Jupiter-mCherry (bottom), together with mRNA encoding ALFA-tagged versions of either WT Spd-2, or Spd-2 mutants that are unable to efficiently recruit Polo (ΔPolo) or AurA (ΔABD1 or ΔABD2). Scale bar = 2μm. (B) Violin plots quantify the fluorescent signal (Median±Quartiles) of the various green- or red-fluorescent fusions (indicated at the top of each plot) that are recruited to the beads when coupled to the different ALFA-tagged Spd-2-fusions (indicated at the base of each plot). Statistical significance was calculated by an Ordinary one-way ANOVA, followed by a Tukey’s Multiple Comparison (****: P<0.0001, ns: not significant).

## Discussion

The prevailing model of mitotic centrosome assembly that has emerged from studies in flies and worms is that Spd-2/SPD-2 recruits Polo/PLK-1, which then phosphorylates Cnn (flies) or SPD-5 (worms) to catalyse their assembly into scaffolds that ultimately recruit all the other PCM proteins. In vertebrates, however, it has long been known that centrosomal CEP192/Spd-2 proteins recruit and activate both PLK1/Polo and AURKA/AurA (Joukov et al., 2014; Meng et al., 2015)—kinases that have several functions at centrosomes and spindle-poles, as well as during mitosis more generally (Asteriti et al., 2015; Joukov and De Nicolo, 2018). It was unclear, however, whether Spd-2/CEP192 proteins also recruit AurA to centrosomes in flies or worms (Joukov and De Nicolo, 2018). We show here that this is the case in flies and that, unexpectedly, this recruitment of AurA stimulates the assembly of a second mitotic PCM scaffold catalysed by TACC phosphorylation. We propose that this Spd-2/CEP192, Polo/PLK1 and AurA/AURKA module is the core driver of mitotic centrosome scaffold assembly in flies (Figure 8A), and likely in many other species— although the molecular mechanisms regulating this module, and the molecular nature of the scaffolds, may vary between species and cell-types.

The centrosomal TACC scaffold we identify here is similar to the liquid-like spindle domain (LISD) originally identified at the poles of acentrosomal mouse oocyte meiotic spindles (So et al., 2019). The assembly of both structures appears to be initiated by the AurA-dependent phosphorylation of TACC/TACC3 (So et al., 2019; Wang et al., 2021; Blengini et al., 2021), which stimulates TACC’s interaction with Clathrin Heavy Chain (CHC) (Lin et al., 2010; Booth et al., 2011; Hood et al., 2013). The LISD was originally postulated to be a unique feature of acentrosomal meiotic spindles because no similar structure was formed on mitotic spindles that were artificially induced to lack centrosomes (So et al., 2019). Our data suggests, however, that centrosomes are required to stimulate TACC scaffold assembly at mitotic spindle poles because they act as a source of Spd-2/CEP192. This would explain why the LISD was not observed on acentrosomal mitotic spindles. Interestingly, CEP192 is present at the poles of acentriolar mouse oocyte spindles (So et al., 2019), but it has not been tested if it is required to recruit AurA and/or to promote LISD assembly.

There has been much interest in the biophysical nature of the mitotic PCM, and in particular in the idea that LLPS might be important for its assembly (Rale et al., 2018; Woodruff, 2018; Raff, 2019; Lee et al., 2021; Woodruff, 2021). Our discovery here that *Drosophila* centrioles generate Polo/Cnn and AurA/TACC scaffolds with solid-like and more liquid-like behaviours, respectively, could be interpreted as at least partial support for the LLPS hypothesis. The disordered N-terminal region of mouse TACC3 can undergo LLPS *in vitro*, leading to the proposal that the LISD forms through LLPS *in vivo* (So et al., 2019), and the N-terminal region of *Drosophila* TACC is also predicted to be disordered—although *C. elegans* TAC-1 appears to lack such a region (Le Bot et al., 2003). The terms solid-like and liquid-like, however, may be intuitively useful for biologists, but they are not very precise (Raff, 2019; Woodruff, 2021). The experiments we describe here simply reveal that molecules in the Cnn-scaffold have little ability to internally rearrange, while molecules in the TACC scaffold re-arrange more-easily. Thus, it is not proven that the TACC scaffold forms via LLPS in *Drosophila*, and we think it plausible that the TACC (and indeed Cnn) scaffold may simply be permeated by the cytoplasm (as water permeates a sponge) rather than phase-separated from the cytoplasm (as oil separates from water). As argued previously, distinguishing between these possibilities *in vivo* is likely to be challenging (Raff, 2019; Musacchio, 2022).

Regardless of their assembly mechanism, it is intriguing that both scaffolds are required for proper centrosome assembly in fly embryos, even though both can recruit PCM and organise MTs independently. The requirement for the solid-like Cnn scaffold is perhaps understandable, as it gives mechanical strength to the PCM (Lucas and Raff, 2007). It is less obvious why centrosomes might require the more liquid-like TACC scaffold. Perhaps the more liquid-like behaviour of the TACC scaffold allows it to expand more easily around the centrioles to form an extended “net” that can efficiently “capture” centriole and centrosome proteins and increase their local concentration. The TACC3 scaffold in human cells (Gergely et al., 2000a), the LISD in mouse oocytes (So et al., 2019), and the TACC scaffold in fly embryos (Barros et al., 2005), all extensively spread around the centrosome/spindle poles, suggesting that the ability to form an extended structure is a conserved feature of the TACC scaffold/LISD. This may be particularly important in the early fly embryo, where centrioles and centrosomes assemble very quickly, yet the cytoplasmic concentration of many key centriole and centrosome proteins is surprisingly low (Steinacker et al., 2022).

An alternative, and not mutually exclusive, possibility is that the existence of more than one mitotic PCM scaffold allows greater flexibility in how client proteins are recruited to centrosomes. We recently showed that mitotic PCM recruitment is a complicated process in fly embryos, and that many centrosome proteins exhibit distinct centrosome-recruitment dynamics (Wong et al., 2024). Moreover, during mitosis Cnn appears to function primarily as a PCM scaffold, whereas it is clear that TACC proteins can perform multiple functions at centrosomes and spindles, some of which are probably not dependent on TACC scaffold assembly (Peset and Vernos, 2008; Thakur et al., 2013; Saatci and Sahin, 2024). Thus, teasing apart the precise function of the TACC scaffold in any given cell type may not be trivial.

An important feature of the scheme depicted in Figure 8A is that it potentially explains why mother centrioles are the dominant sites of mitotic PCM assembly, as they provide a source of Spd-2/CEP192. Our bead-reconstitution experiments provide strong support for this hypothesis. Simply concentrating Spd-2 on the bead surface triggers several aspects of mitotic PCM assembly and MT organisation, whereas similarly concentrating Cnn or TACC molecules does not. Interestingly, the N-terminal 1000aa of *Xenopus* CEP192 can also organise MTs when recruited to synthetic beads in *Xenopus* mitotic extracts (Joukov et al., 2014). The ability of Spd-2/CEP192 to recruit AurA/AURKA and/or Polo/PLK1 is required for bead-dependent MT organisation in both fly embryos and *Xenopus* extracts. Clearly there is an intimate and complicated reciprocal relationship between AurA/AURKA and Polo/PLK1 kinase activation (Asteriti et al., 2015; Joukov and De Nicolo, 2018), and more work is required to understand how this occurs in the context of their Spd-2/CEP192-dependent recruitment to centrosomes.

Finally, our bead reconstitution experiments potentially shed light on the importance of the outward flux of Spd-2 from centrioles. We previously inferred this flux from FRAP experiments, and speculated that it might be important for setting mitotic centrosome size—as the size of the PCM-scaffold would be limited by the amount of Spd-2 that the centrioles can generate (Conduit et al., 2015; Raff, 2019). Here, we directly demonstrate the outward flux of Spd-2 from the centrioles. This flux, however, presumably does not occur on the bead surface (as Spd-2-GFP molecules are recruited via a high-affinity interaction with the GFP-Nanobody), and the Cnn scaffold remains tightly associated with the bead surface. In contrast, the TACC scaffold can extensively extend outwards around the bead. Thus, the outward flux of Spd-2 molecules may be important for the expansion of the solid-like Cnn scaffold, but not for the expansion of the more liquid-like TACC scaffold. It is not clear if a similar Spd-2 flux exists in other species—and FRAP experiments provided no evidence for it in *C. elegans* embryos (Cabral et al., 2019)—but our observations demonstrate conclusively that it occurs in flies, where is appears to be of functional importance.

## Acknowledgements

We are grateful to members of the Raff Laboratory for advice, discussion and for critically reading the manuscript. The research was funded by: a Wellcome Trust Senior Investigator Award (215523) (JWR, supporting AW, SS, NM and TS); a Cancer Research UK Oxford Centre Prize DPhil Studentship (C5255/A23225), a Balliol Jason Hu Scholarship, a Clarendon Scholarship, a Max Planck Croucher Postdoctoral Fellowship and a Junior Research Fellowship in Medical Sciences from the Wadham College (SSW): a Marie-Sklodowska-Curie Individual Fellowship 892857 – CENTROMD, from the European Commission and by LTF 29-2019 from the European Molecular Biology Organisation (JMM); an Overseas Project for Post Graduate Research from the National Science and Technology Council, Taiwan, ROC #111-2917-I-564-012 (CCC) and #112-2917-I-002-026 (MP).

## Author contributions

**SSW**: Conceptualisation; Data curation; Software; Formal analysis; Investigation; Methodology; Writing—review and editing. **JMM**: Conceptualisation; Data curation; Formal analysis; Investigation; Methodology; Writing—review and editing. **CCC**: Conceptualisation; Data curation; Formal analysis; Investigation; Methodology; Writing—review and editing. **MP, NM, TS, SS, AW**: Data curation; Formal analysis; Investigation; Writing—review and editing. **JWR:** Conceptualisation; Formal analysis; Funding acquisition; Project administration; Writing—review and editing; Writing— original draft.

## Rights Retention Statement

This research was funded in whole or in part by Wellcome (215523). For the purpose of Open Access, the author has applied a CC BY public copyright licence to any Author Accepted Manuscript (AAM) version arising from this submission.

## Materials and Methods

### Drosophila melanogaster stocks and husbandry

The *Drosophila* stocks used, generated and/or tested in this study are listed in Table 1. The precise stocks used in each experiment (and the relevant Figure) are listed in Table 2. Flies were maintained on *Drosophila* culture medium (0.68% agar, 2.5% yeast extract, 6.25% cornmeal, 3.75% molasses, 0.42% propionic acid, 0.14% tegosept, and 0.7% ethanol) in 8cm x 2.5cm plastic vials or 0.25-pint plastic bottles. For microscopy and immunoblot experiments, flies were placed in embryo collection cages on fruit juice plates (see below) with a drop of yeast paste. Fly handling were performed as previously described (Roberts, 1998).

**Table 1:** Drosophila stocks used in this study.

| Allele | Source |
| --- | --- |
| cnn<sup>f04547</sup> | Exelixis stock no. f04547, Exelixis Stock Centre (Harvard Medical School, Boston, MA). |
| cnn<sup>HK21</sup> | (Megraw et al., 1999; Vaizel-Ohayon and Schejter, 1999) |
| GFP-Cnn | (Tovey et al., 2020); CRISPR Knock-in fast folded GFP |
| Ubq-GFP-TACC | (Barros et al., 2005) |
| Ubq-NG-Cnn | (Wong et al., 2022) |
| tacc<sup>stella592</sup> | (Lee et al., 2001) |
| aur<sup>1</sup> | (Glover et al., 1995) |
| plp<sup>2172</sup> | (Spradling et al., 1999) |
| Spd-2<sup>z35711</sup> | (Giansanti et al., 2008b) |
| Plk4<sup>Aa74</sup> | (Aydogan et al., 2018) |
| Ubq-RFP-Cnn | (Conduit & Raff, 2010) |
| Ubq-GFP-TACC-S863L | (Barros et al., 2005) |
| Ubq-NG-TACC | Generated in this study |
| Ubq-NG-TACC-S863L | Generated in this study |
| tacc<sup>59</sup> | (Tang et al., 2020); Gift from Fengwei Yu |
| tacc<sup>74</sup> | (Tang et al., 2020); Gift from Fengwei Yu |
| Ubq-Aurora A-GFP | (Lucas and Raff, 2007) |
| Ubq-Spd-2-RFP | (Conduit et al., 2014c) |
| Ubq-Spd-2-SNAP | Generated in this study |
| Asl-NG | (Steinacker et al., 2022) |
| Ubq-Spd-2-NG | (Wong et al, 2022) |
| Ubq-Msps-GFP | (Lee et al., 2001) |
| Ubq-Klp10A-GFP | Generated in this study |
| ncd-γ-tubulin-37c-GFP | (Hallen et al., 2008) |
| UAS-CHC-GFP | (Li et al., 2010) |
| Ubq-GFP-Cnn-ΔLZ | (Feng et al., 2017) |
| Ubq-GFP-Cnn-ΔCM2 | (Feng et al., 2017) |
| (pSas-6)-mNG | (Steinacker et al., 2022) mNeonGreen expression driven by the Sas-6's promoter |
| NG-Sas-6 | (Steinacker et al., 2022) |
| Ana2-NG | (Steinacker et al., 2022) |
| Spd-NG | (Wong et al., 2022) |
| NG-Cnn | (Wong et al., 2022) |
| Polo-TRAP-GFP | (Buszczak et al., 2007); appears to not be fully functional and is only viable as a heterozygote. |
| Ubq-RFP-Cnn | (Lucas and Raff, 2007b) |
| Ubq-mCherry-TACC | Generated in this study |
| Ubq-mCherry-TACC-S863L | Generated in this study |
| Jupiter-mCherry | (Karpova et al., 2006) |

**Table 2:** Drosophila stocks used in specific experiments.

| Genotype | Purpose | Figure |
| --- | --- | --- |
| GFP-Cnn | Comparing the characteristics of Cnn and TACC in cycling or colchicine-treated embryos | 1A-B, 2A-D |
| Ubq-GFP-TACC/+ | Comparing the characteristics of Cnn and TACC in cycling or colchicine-treated embryos<br>Comparing the properties of TACC in the presence or absence of endogenous Cnn | 1A-B, 1D, 2A-D, S1, S3A |
| Ubq-GFP-TACC-S863L/+ | Comparing the expression levels to Ubq-GFP-TACC | S3A |
| Ubq-GFP-TACC, Ubq-RFP-Cnn | Showing two scaffolds together in the same embryos | 1C |
| cnn<sup>f04547</sup> / cnn<sup>HK21</sup> ; Ubq-GFP-TACC / + | Comparing the properties of TACC in the presence or absence of endogenous Cnn<br>Comparing the size of TACC in various genetic background in the absence of endogenous Cnn | 1D, 3, S1 |
| cnn<sup>f04547</sup> / cnn<sup>HK21</sup> ; Ubq-GFP-TACC / tacc<sup>stella592</sup> | Comparing the size of TACC in various genetic background in the absence of endogenous Cnn | 3 |
| cnn<sup>f04547</sup> / cnn<sup>HK21</sup> ; Ubq-GFP-TACC / aur<sup>1</sup> |  |  |
| chc<sup>4</sup>/+ ; cnn<sup>f04547</sup> / cnn<sup>HK21</sup> ; Ubq-GFP-TACC / + |  |  |
| cnn<sup>f04547</sup> / cnn<sup>HK21</sup> ; Ubq-GFP-TACC / plp<sup>2172</sup> |  |  |
| cnn<sup>f04547</sup> / cnn<sup>HK21</sup> ; Ubq-GFP-TACC / Plk4<sup>Aa74</sup> |  |  |
| cnn<sup>f04547</sup> / cnn<sup>HK21</sup> ; Ubq-GFP-TACC / Spd-2<sup>z35711</sup> |  |  |
| cnn<sup>f04547</sup> , RFP-Cnn / +; Ubq-GFP-TACC, tacc<sup>stella592</sup> / + | Comparing the size of WT-TACC and non-phosphorylatable TACC-S863L mutants relative to the size of Cnn scaffold | 4A, S3B |
| cnn<sup>f04547</sup> , RFP-Cnn / +; Ubq-GFP-TACC-S863L, tacc<sup>stella592</sup> / + |  |  |
| Ubq-NG-TACC, tacc<sup>59</sup> / tacc<sup>74</sup> | Comparing the recovery of WT-TACC and non-phosphorylatable TACC-S863L mutants in the absence of endogenous TACC | 4B, S3B |
| Ubq-NG-TACC-S863L, tacc<sup>59</sup> / tacc<sup>74</sup> |  |  |
| Ubq-Aurora A-GFP | Comparing the recruitment of Aurora A-GFP in embryos injected with different Spd-2 fused with mScarlet-I3 | 5B |
| Ubq-Aurora A-GFP / Ubq-Spd-2-RFP | Comparing the recovery of Spd-2 and Aurora A in the same embryos | 6A-B |
| Ubq-Spd2-SNAP / +; Spd2-NG / Spd-2<sup>z35711</sup> | Selective labelling of single molecules, Spd-2 single molecule behaviour | 6C |
| Ubq-Spd2-SNAP / +; Asl-NG / Spd-2<sup>z35711</sup> | Measuring the distance between single Spd-2 molecules and the centriole labelled by Asl-NG | 6D-E |
| Ubq-Spd-2-NG / + | Comparing the relative enrichment of various centrosomal proteins with or without Cnn scaffold | 7A |
| cnn<sup>f04547</sup> / cnn<sup>HK21</sup> ; Ubq-Spd-2-NG / + |  |  |
| Ubq-Aurora A-GFP / + |  |  |
| cnn<sup>f04547</sup> / cnn<sup>HK21</sup> ; Ubq-Aurora A-GFP / + |  |  |
| ncd-γ-tubulin-37c-GFP / + |  |  |
| cnn<sup>f04547</sup> / cnn<sup>HK21</sup> ; ncd-γ-tubulin-37c-GFP / + |  |  |
| Ubq-Msps-GFP / + |  |  |
| cnn<sup>f04547</sup> / cnn<sup>HK21</sup> ; Ubq-Msps-GFP / + |  |  |
| Ubq-Klp10A-GFP / + |  |  |
| cnn<sup>f04547</sup> / cnn<sup>HK21</sup> ; Ubq-Klp10A-GFP / + |  |  |
| V32a-Gal4 / + ; UAS-CHC-GFP / + |  |  |
| cnn<sup>f04547</sup>, V32a-Gal4 / cnn<sup>HK21</sup> ; UAS-CHC-GFP / + |  |  |
| cnn<sup>f04547</sup>, Ubq-GFP-Cnn-ΔLZ / + | Showing Cnn-mutants that do not form scaffold can be concentrate to the TACC scaffold | 7B |
| cnn<sup>f04547</sup>, Ubq-GFP-Cnn-ΔLZ / cnn<sup>HK21</sup> |  |  |
| cnn<sup>f04547</sup>, Ubq-GFP-Cnn-ΔCM2 / + |  |  |
| cnn<sup>f04547</sup>, Ubq-GFP-Cnn-ΔCM2 / cnn<sup>HK21</sup> |  |  |
| cnn<sup>HK21</sup> / + ; (pSas-6)-mNG / Ubq-RFP-TACC | Comparing the concentration and diffusion coefficient of various centrosomal proteins inside and outside of TACC scaffold | 7C, S5 |
| cnn<sup>f04547</sup> / cnn<sup>HK21</sup> ; (pSas-6)-mNG / Ubq-RFP-TACC |  |  |
| cnn<sup>HK21</sup> / + ; NG-Sas-6/ Ubq-RFP-TACC |  |  |
| cnn<sup>f04547</sup> / cnn<sup>HK21</sup> ; NG-Sas-6/ Ubq-RFP-TACC |  |  |
| cnn<sup>f04547</sup>, Ana2-NG / + ; Ubq-RFP-TACC / + |  |  |
| cnn<sup>f04547</sup>, Ana2-NG / cnn<sup>HK21</sup> ; Ubq-RFP-TACC / + |  |  |
| Spd-2-NG / Ubq-RFP-TACC |  |  |
| NG-Cnn / + ; Ubq-RFP-TACC / + |  |  |
| cnn<sup>f04547</sup>, RFP-Cnn / + | Comparing the localisation of RFP-Cnn, mCherry-TACC or mCherry-Jupiter on synthetic beads coated with α-GFP nanobody in cycling embryos co-injected various GFP-fused centrosomal proteins | 8 |
| Ubq-mCherry-TACC |  |  |
| Jupiter-mCherry |  |  |
| cnn<sup>f04547</sup>, RFP-Cnn / Polo-TRAP-GFP | Comparing the localisation of RFP-Cnn, mCherry-TACC or mCherry-Jupiter co- | 9 |
| Ubq-mCherry-TACC / Aurora A-GFP | expressed with Polo-GFP or Aurora-GFP on synthetic beads coated with α-ALFA nanobody in cycling embryos co-injected various Spd-2 mutants fused with ALFA-tag | |
| Jupiter-mCherry / Polo-TRAP-GFP |  |  |
| Jupiter-mCherry / Aurora A-GFP |  |  |
| cnn<sup>f04547</sup>, Spd-2-RFP / cnn<sup>HK21</sup> ; Ubq-GFP-TACC / + | Observing GFP-TACC nucleation and dissipation from centrioles | S4 |

### Transgenic fly line generation

Transgenic fly lines were generated via random P-element insertion (injected, mapped, and balanced by ‘The University of Cambridge Department of Genetics Fly Facility’). For transgene selection, the *w^+^*gene marker was included in the transformation vectors and constructs were injected into the *w^1118^* genetic background.

### Molecular biology

To generate NG-TACC transgenic flies, cDNA fragment encoding TACC was amplified using primers s1 and s2 (Table 3) containing the attB sites and transferred to a pDONR plasmid backbone using a BP clonase (Gateway technology, ThermoFisher Scientific) to generate a pDONR-TACC. For non-phosphorylatable mutant TACC-S863L, primers s3 and s4 (Table 3) were used to create a single amino acid mutation on the pDONR-TACC. Flipping TACC or TACC-S863L from pDONR vector to pUbq-mNeonGreen (NG) or pUbq-mCherry destination vector using a LR clonase (Gateway technology, ThermoFisher Scientific), a N-terminally fused pUbq-NG-TACC or pUbq-NG-TACC-S863L or pUbq-mCherry-TACC or pUbq-mCherry-TACC-S863L plasmids were generated. We note that Serine originally designated as Ser863 in this previous paper is Ser900 in the 1227aa long TACC-PA isoform listed in FlyBase, which is the isoform we use in the experiments reported here. To generate Ubq-Klp10A-GFP an entry vector containing Klp10A without stop codon (pENTR4-Klp10A_LD29208 cDNA) was obtained (Radford et al., 2012). Klp10A was then introduced into pUbq-mGFP destination vector using LR clonase (Gateway technology, ThermoFisher Scientific).

**Table 3:**
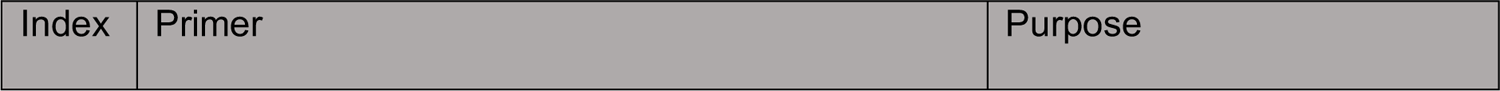
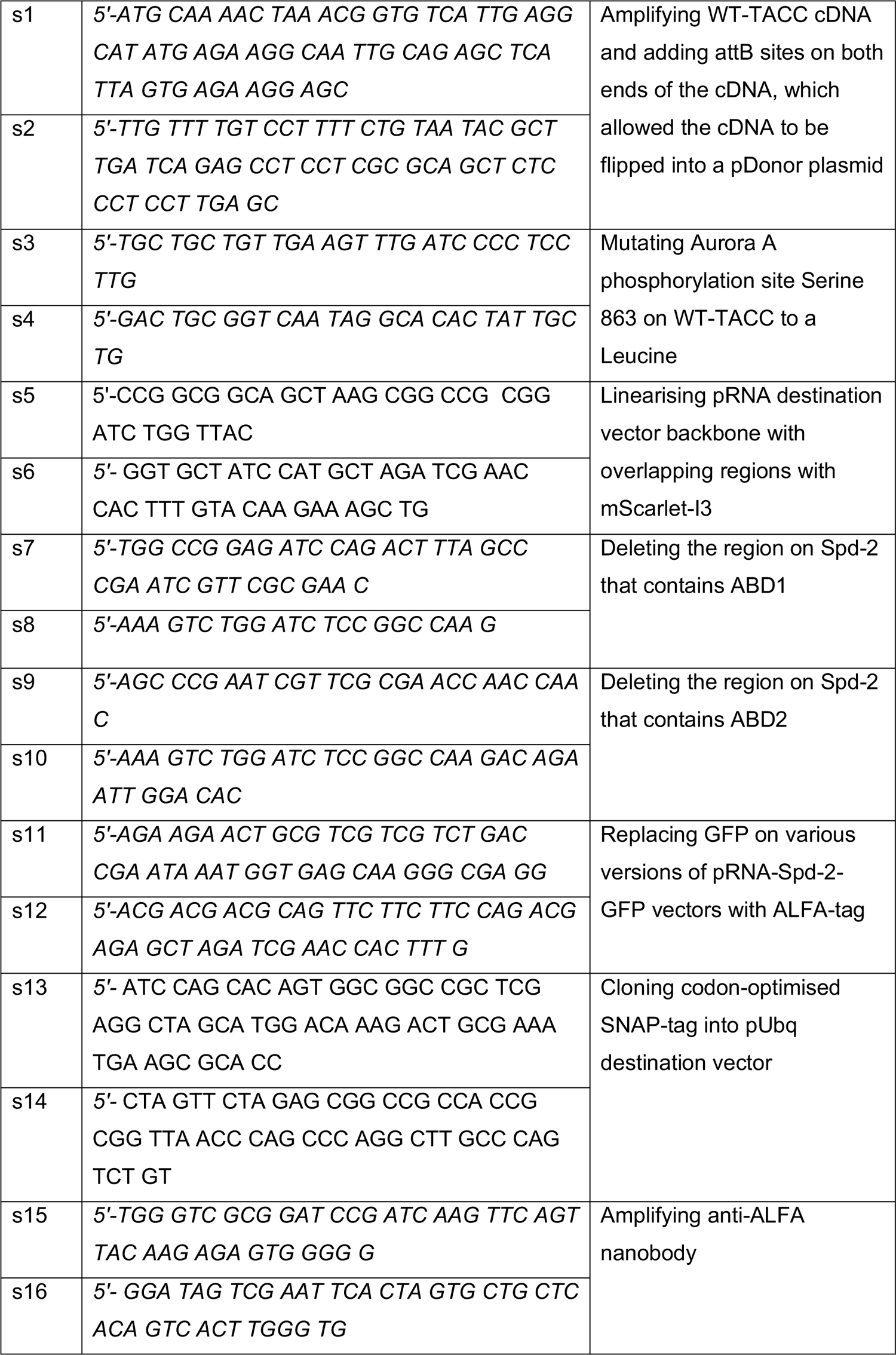
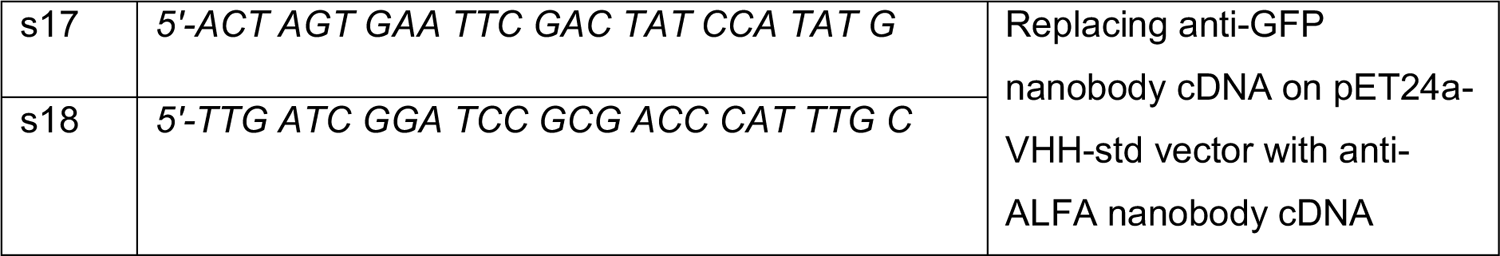
Primers used in this study.

**Table 4:** Plasmids used in this study.

| Index | Plasmid | Purpose | Source |
| --- | --- | --- | --- |
| p1 | pDonor-TACC-WT | Donating WT-TACC to various pDestination vectors | Generated in Raff Lab |
| p2 | pDonor-TACC-S863L | Donating TACC-S863L to various pDestination vectors | Generated in Raff Lab |
| p3 | pUbq-NG-NT destination | Adding NG to the N-terminus of donor cDNA for transgenic fly generation | Generated in Raff Lab |
| p4 | pUbq-mCherry-NT destination | Adding mCherry to the N-terminus of donor cDNA for transgenic fly generation | Generated in Raff Lab |
| p5 | pUbq-NG-TACC-WT | Generating transgenic fly line expressing N-terminally NG-fused WT-TACC | Generated in Raff Lab |
| p6 | pUbq-NG-TACC-S863L | Generating transgenic fly line expressing N-terminally NG-fused TACC-S863L | Generated in Raff Lab |
| p7 | pUbq-mCherry-TACC-WT | Generating transgenic fly line expressing N-terminally mCherry-fused WT-TACC | Generated in Raff Lab |
| p8 | pUbq-GFP-CT destination | Adding GFP to the C-terminus of donor cDNA for transgenic fly generation | Generated in Raff Lab |
| p9 | pUbq-Klp10A-GFP | Generating transgenic fly line expressing C-terminally GFP-fused Klp10A | Generated in Raff Lab |
| p10 | pRNA-GFP-CT destination | Adding GFP to the C-terminus of donor cDNA for mRNA synthesis | Generated in Raff Lab |
| p11 | pDonor-Spd-2-WT | Donating Spd-2-WT to various pDestination vectors | Generated in Raff Lab |
| p12 | pRNA-Spd-2-GFP | Synthesizing mRNA encoding C-terminally GFP-fused WT-Spd-2 | Generated in Raff Lab |
| p13 | pRNA-GFP-NT destination | Adding GFP to the N-terminus of donor cDNA for mRNA synthesis | Generated in Raff Lab |
| p14 | pDonor-Cnn-WT | Donating Cnn-WT to various pDestination vectors | Generated in Raff Lab |
| p15 | pRNA-GFP-Cnn | Synthesizing mRNA encoding N-terminally GFP-fused WT-Cnn | Generated in Raff Lab |
| p16 | pRNA-GFP-TACC | Synthesizing mRNA encoding N-terminally GFP-fused WT-TACC | Generated in Raff Lab |
| p17 | pDRESS_mTurquoise2_spatial-linker-P2A_mScarlet-I3 | mScarlet-I3 containing plasmid | Addgene #189755 |
| p18 | pRNA-mScarlet-I3 Destination | Adding mScarlet-I3 to the C-terminus of donor cDNA for mRNA synthesis | Generated in Raff Lab |
| p19 | pDonor-Spd-2ΔPolo | Donating Spd-2ΔPolo to various pDestination vectors | Generated in Raff Lab |
| p20 | pRNA-Spd-2ΔPolo-mScarlet-I3 | Synthesizing mRNA encoding C-terminally mScarlet-I3-fused Spd-2ΔPolo | Generated in Raff Lab |
| p21 | pRNA-Spd-2ΔABD1-mScarlet-I3 | Synthesizing mRNA encoding C-terminally mScarlet-I3-fused Spd-2ΔABD1 | Generated in Raff Lab |
| p22 | pRNA-Spd-2ΔABD2-mScarlet-I3 | Synthesizing mRNA encoding C-terminally mScarlet-I3-fused Spd-2ΔABD2 | Generated in Raff Lab |
| p23 | pRNA-Spd-2-ΔABD1-ALFA | Synthesizing mRNA encoding C-terminally ALFA-fused Spd-2ΔABD1 | Generated in Raff Lab |
| p24 | pRNA-Spd-2-ΔABD2-ALFA | Synthesizing mRNA encoding C-terminally ALFA-fused Spd-2ΔABD2 | Generated in Raff Lab |
| p25 | pUbq-SNAP Destination | Adding SNAP to the C-terminus of donor cDNA for transgenic fly generation | Generated in Raff Lab |
| p26 | pUbq-Spd-2-SNAP | Generating transgenic fly line expressing C-terminally SNAP-fused Spd-2 | Generated in Raff Lab |
| p27 | cfSGFP2-anti-AlfaTag nanobody | anti-ALFA nanobody containing plasmid | Addgene #171818 |
| p28 | pET24a-VHH-std | Expressing anti-GFP nanobody | Addgene #109417 |
| p29 | pET24a-anti-ALFA-std | Expressing anti-ALFA nanobody | Generated in Raff Lab |

To generate Spd-2-GFP, GFP-Cnn, and GFP-TACC constructs for in vitro mRNA synthesis, pDONR vectors containing Spd-2, Cnn or TACC CDS were recombined with a destination pRNA vector containing a GFP CDS at either the N- or C-terminus using Gateway technology (ThermoFisher Scientific).

For various Spd-2 variants fused to mScarlet-I3 (Gadella et al., 2023), a mScarlet-I3 expression vector was purchase from Addgene (#189755), and later optimised to Drosophila codon usage by Genewiz. The pRNA destination backbone was linearised by primers s5 and s6 (Table 3) and assembled with the mScarlet-I3 fragment using a NEBuilder® HiFi DNA Assembly kit (NEB) to create a pRNA-mScarlet-I3 destination vector. pDonor vector containing Spd-2 cDNA was flipped into pRNA-mScarlet-I3 destination vector using LR clonase (Gateway technology, ThermoFisher Scientific) to generate pRNA-Spd-2-mScarlet-I3. Similarly, a pDonor containing Spd-2-ΔPolo (previously known as Spd-2-ALL) was flipped to generate a pRNA-Spd-2-ΔPolo-mScarlet-I3. For Spd-2-ΔABD1 and Spd-2-ΔABD2, the targeted sequences in pRNA-Spd-2-mScarlet-I3 and pRNA-Spd-2-GFP were removed by site-directed mutagenesis using either primers s7 and s8, or primers s9 and s10 (Table 3) to generate linearised pRNA-Spd-2-ΔABD1-mScarlet-I3, pRNA-Spd-2-ΔABD1-GFP, pRNA-Spd-2-ΔABD2-mScarlet-I3 and pRNA-Spd-2-ΔABD2-GFP, respectively. pRNA-Spd-2-ΔABD1-GFP and pRNA-Spd-2-ΔABD2-GFP were linearised by primers s11 and s12 (Table 3) to introduce an ALFA sequence followed by a stop codon in the C-terminus of Spd-2, which can be efficiently recognised by anti-ALFA nanobody (Götzke et al., 2019).

For Spd-2-SNAP, a pDONR vector containing a full length Spd-2 cDNA encoding the was used. pSNAPf vector containing SNAP-tag was purchased from NEB (N9183S) and optimised to Drosophila codon suing Genewiz (USA) which was subcloned to make a pUbq-SNAP destination vector using primers s13 and s14 (Table 3), which contain regions from a pUbq destination vector. The amplified fragment containing SNAP was assembled with an amplified pUbq destination backbone using a NEBuilder® HiFi DNA Assembly kit (NEB) to create a pUbq-SNAP destination vector. Spd-2 from a pDONR vector was introduced into pUbq-SNAP destination vector using LR clonase (Gateway technology, ThermoFisher Scientific).

To generate expression plasmid for anti-ALFA nanobody, anti-ALFA nanobody fragment was amplified using primers s15 and s16 (Table 3) from its expression vector (Addgene #189755). A pET24a-VHH-std vector (Addgene #109417) was linearised by primers s17 and s18 (Table 3). that overlapped the sequence of anti-ALFA nanobody. These fragments were assembled using a NEBuilder® HiFi DNA Assembly kit (NEB) to create a pET24a-anti-ALFA-std expression vector. The list of primers and plasmids used or generated in this study are listed in Table 3 and 4.

### Nanobody-coated beads preparation

0.1 mg of Dynabeads™ MyOne™ Streptavidin C1 (65001, Invitrogen) were washed 3X in PBS, supplemented with 0.01% Triton X-100 (T9284, Sigma-Aldrich) and 0.1% Bovine Serum Albumin (A7906, Sigma-Aldrich) (PBSTB). The beads were incubated with either anti-GFP or anti-ALFA nanobody (4.4 µg/ml—prepared as described below) in PBSTB for a minimum of 30min at room temperature. The beads were then washed 3X in PBSTB and stored at a concentration of 3mg/ml at 4°C until further use (for maximum efficiency, the beads were used within 1 week after preparation). For embryo injection, the nanobody-coated beads were diluted with either mRNA solution or water to achieve a final concentration of 1mg/ml.

### Nanobody Expression and Purification

Recombinant anti-GFP (Buser et al., 2018) or anti-ALFA-tag (Götzke et al., 2019) nanobody was co-transformed with pET-21d-myc-BirA (to biotinylate the Nanobody) into Rosetta (DE3) bacterial cells. Cells were grown on a shaking incubator using the appropriate antibiotics and 200μM dbiotin in LB broth at 37°C until an OD600 of 0.6-0.7 was reached. Protein expression was then induced with 1mM Isopropyl β-d-1-thiogalactopyranoside (IPTG) and cells were grown overnight at 18°C. Cells were pelleted, washed, resuspended in binding PBS buffer supplemented with Complete EDTA-free Protease inhibitor Cocktail (Roche) and lysed using an Emulsiflex-C5 homogeniser (Avestin). The soluble protein fraction was then purified using a HisTrap HP prepacked column (Cytiva). An extra purification step of Size exclusion chromatography was carried out using an S75 16/600 column (GE healthcare) equilibrated against a PBS buffer pH 7.5. Purified anti-GFP or anti-ALFA-tag nanobodies were stored in Liquid Nitrogen at ∼1mg/ml or ∼5.5mg/ml respectively.

### mRNA synthesis

For *in vitro* transcription, the constructs were digested and linearised by AscI (R0558S, NEB), precipitated with 10 mM sodium acetate and 7 mM EDTA in 66% ethanol overnight at −20 °C. Precipitated DNA was washed with 70% Ethanol, dried and subsequently dissolved in DEPC-treated water (AM9906, Ambion). Around 1.6-3.2μg of digested DNA was used to synthesise mRNA with the T3 mMESSAGE mMACHINE kit (AM1348, ThermoFisher Scientific). The mRNA product was purified with the RNeasy MinElute Cleanup Kit (74204, Qiagen). All RNAs were stored at −70°C. The final concentrations of RNA used in various experiments are described in the next section.

### Embryo collection and injection

Embryos were collected from plates (40% cranberry-raspberry juice, 2% sucrose, and 1.8% agar) supplemented with fresh yeast suspension. For live-imaging experiments, embryos were collected for 1h at 25°C, and aged at 25°C for 45–60 min. Embryos were dechorionated by hand, mounted on a strip of glue on either a 35-mm glass-bottom Petri dish with 14 mm micro-well (MatTek) or high precision 35-mm, high glass bottom μ-dishes (ibidi; for FCS experiments), and desiccated for 1 min at 25°C before covering with Voltalef oil (H10S PCTFE, Arkema) to avoid further desiccation.

For colchicine injection, embryos were desiccated for 5-10min (depending on the ambient conditions) before covering with Voltalef oil and injecting 100μg/ml colchicine (diluted in Schneider’s medium from a 1mg/ml stock in DMSO). Embryos were imaged on the PerkinElmer spinning disk system described below.

For single molecule experiments, embryos were aged at 25°C for 30 min, and desiccated for 5min. After covering with Voltalef oil, they were injected with SNAP dye JF549 (GA1110, Promega), dissolved in DMSO (Grimm et al., 2015) at concentrations ranging from 1-1000 nM and incubated for a further 30 min at 25°C prior to imaging on the Andor DragonFly 505 (Oxford Instruments) spinning disk system described below.

For synthetic bead injection, nanobody-coated beads were mixed with mRNA or water to achieve a final concentration of 1 mg/ml beads and 150μg/ml mRNA. Embryos were collected for 30 min at 25°C, dechorionated, and desiccated for 5 min. After covering with Voltalef oil, embryos were injected with the nanobody-coated beads and mRNA mix and incubated for a further 30 min at 25°C prior to imaging on the Andor DragonFly 505 (Oxford Instruments) spinning disk system described below.

For mRNA only injection, mRNA was diluted to a final. concentration of 2mg/ml. Embryos were collected for 30 min at 25°C, dechorionated, and desiccated for 5 min. After covering with Voltalef oil, embryos were injected with the nanobody-coated beads and mRNA mix and incubated for a further 1 hour at 22°C prior to imaging on the Andor DragonFly 505 (Oxford Instruments) spinning disk system described below.

### Spinning disk confocal microscopy

Images of living embryos were acquired at 23°C using a PerkinElmer ERS spinning disk confocal system mounted on a Zeiss Axiovet 200M microscope using Volocity software (PerkinElmer). A 63X, 1.4NA oil objective was used for all acquisition. The oil objective was covered with an immersion oil (ImmersolT 518 F, Carl Zeiss) with a refractive index of 1.518 to minimize spherical aberration. The detector used was a charge-coupled device (CCD) camera (Orca ER, Hamamatsu Photonics, 15-bit), with a gain of 200 V. The system was equipped with 405nm, 488nm, 561nm, and 642 solid-state lasers (Oxxius S.A.). The microscope was operated using a Volocity software. All red/green fluorescently tagged samples were acquired using UltraVIEW ERS ‘Emission Discrimination’ setting. The emission filters used were a green long-pass 520nm emission filter and a red long-pass 620nm emission filter. For dual channel imaging, the red channel was imaged before the green channel in every slice in a z-stacks. 0.5-μm z-sections were acquired, with the number of sections, time step, laser power, and exposure depending on the experiment. For Fluorescent Recovery after Photobleaching (FRAP) experiments, multiple circular regions of interests (ROI) with a diameter of 3.5-4μm were created around different centrosomes. 488nm and 521nm lasers with 50% laser power of 20 iterations were used to FRAP each sample. In some samples, different centrosomes were bleached at different timepoints in the same embryos. 0.5-μm z-sections were acquired, with the number of sections, time step, laser power, and exposure depending on the experiment.

For single molecule, synthetic bead and mRNA injection experiments, embryos were imaged on an Andor Dragonfly 505 (Oxford Instruments) spinning disk confocal microscope (40μm pinhole size), which was mounted on a Leica DMi8 stand, using Fusion software. 561 and 488nm solid-state diode lasers were used to image JF549 and mNG, respectively, using a HCPL APO 63×/1.40 oil immersion objective and an Andor iXon Ultra 888 EMCCD camera. Stacks consisting of 8 slices with a z spacing of 0.5μm were acquired every 10 seconds for 30 minutes total duration for single molecule tracking. For synthetic bead analysis, a single stack consisting of 41 slices with a z spacing of 0.5 μm were collected for each embryo. For mRNA injection experiments, an Andor Sona CMOS camera was used image embryos injected with various Spd-2 variants. A single stack consisting of 17 slices with a z spacing of 0.5μm were collected for each embryo.

### Super-resolution spinning disk confocal microscopy

Super-resolution imaging was performed using a SoRA disk (Yokagawa), a 3.2X magnification lens (Olympus), and a photometrics BSI camera (95% QE – 6.5μm pixels) mounted on an Olympus IX83 microscope equipped with a 60X, 1.3NA silicon immersion lens. The oil objective was covered with an immersion oil (ImmersolT 518 F, Carl Zeiss) with a refractive index of 1.518 to minimize spherical aberration. Samples were excited with QBIS 488 or 561nm laser (Coherant) with 525/50 or 617/75 nm emission bandpass filters, respectively. A ‘Quad’ dichoric of 405/488/561/640 nm was used. At least 6 z-sections with each of 0.5 μm-thickness were used for each image. Laser power and exposure were adjusted according to the experiment.

### Fluorescence correlation spectroscopy (FCS)

Point FCS measurements were performed and analysed as previously described (Aydogan et al., 2020). All measurements were conducted on a confocal Zeiss LSM 880 (Argon laser excitation at 488 nm and GaAsP photon-counting detector [491–544 nm detector range]) with Zen Black Software. A C-Apochromat 40×/1.2 W objective and a pinhole setting of 1AU was used, and spherical aberrations were corrected for on the correction collar of the objective at the beginning of each experimental day by maximizing the FCS-derived CPM value of a fluorescent dye solution. The effective volume Veff of this system was previously estimated to be ∼0.25fL (Steinacker et al., 2022). Measurements were conducted with a laser power of 6.31μW and no photobleaching was observed for any protein. The temperature of the microscope was kept between 25.0 and 26.0°C using the Zeiss inbuilt heating unit XL. For experimental FCS recordings, consecutive cytoplasmic measurements were made 6× for 4s each at the centrosomal plane of the embryo. For measurements within the PCM, a snapshot of the centrosomes in the embryo was taken prior to initiating the FCS measurements, and at the end of the FCS measurements, and the data discarded if the centrosome had moved away from the measurement point. In addition, any erratic autocorrelation functions from the cytoplasmic measurements (usually generated when a centrosome or yolk granule moved into the point of measurement) were also discarded. All remaining curves were then fitted with eight different diffusion models in the FoCuS-point software, including one or two diffusing species with no dark state of the fluorophore, one dark state of the fluorophore (either triplet or blinking state), or two dark states of the fluorophore (triplet and blinking state; Waithe et al., 2016).

### Image data analysis

For centrosome analysis, raw images were z-projected using the maximum intensity projection function, and the background was estimated and corrected using an uneven illumination background correction using a custom Python script. Centrosomes were detected and located by a Crocker-Grier centroid-finding algorithm (Crocker and Grier, 1996) available in a Python library TrackPy (Allan et al., 2016). The signal-to-background intensity threshold was calculated by an Otsu algorithm (Otsu, 1979). The sum intensity and area of each centrosome were calculated from the segmented region based on the Otsu threshold.

FRAP data was initially analysed using Fiji. Raw time-series images were corrected for photobleaching using the exponential decay function, z-projected using the maximum intensity projection function, and the background was estimated and corrected using an uneven illumination background correction (Soille, 2004). Photobleached centrosomes were manually tracked using a software package developed for mobile robotics (Cartucho et al., 2018). A custom Python script was used to extract the fluorescence intensities at tracked centrosomes as they changed over time in each individual embryo, as previously described (Wong et al., 2022).

Synthetic bead fluorescence intensity data was analysed using Fiji. Raw single-frame images were z-projected using the maximum intensity projection function, and background was corrected using an uneven illumination background correction. GFP beads were auto-selected and their intensities were calculated by the TrackMate plugin (Tinevez et al., 2016) with an estimated object diameter of 2.7 microns. The fluorescence mean intensity of each bead was measured and analysed using scripts generated by ChatGPT4, before visualised using GraphPad Prism.

The single molecule Spd-2-NG data was analysed using Fiji (ImageJ2 Ver.2.3.0/1.53q) Fiji software (Schindelin et al, 2012). Hyperstacks corresponding to time-lapse videos of embryos were maximum intensity projected onto a single Z-plane and bleach corrected. Trackmate (Tinevez et al., 2016) was used to detect and track single particles of JF549 covalently bound to SNAP-tagged Spd-2. Single molecules were defined with an estimated object diameter of 0.5 µm in the orange channel with a LoG detector and were tracked over time, using a simple LAP tracker, with linking max distance 2.0 μm, gap-closing max distance 2.0 μm and gap-closing max frame gap of 1. Each tracked particle was inspected for co-localisation with the mother centriole— recognised with the Asl-NG marker (Novak et al., 2014). The following rules were used as criteria to select single molecule-binding events: 1) Only embryos with sparse labelling (<5% centrosomes labelled at any one timepoint) were analysed; 2) Incomplete tracks, in which we did not detect the molecule entering and/or leaving the centrosome, were excluded; 3) Tracks in which molecules were only observed at the centrosome for one timepoint were excluded; 4) We included tracks where molecules disappeared for a single timepoint, but then reappeared in the next, as single fluorophores can blink into temporary dark states (Dickson et al., 1997); 5) The occasional centrosomes where more than one binding event occurred at the same time were excluded. Mother centrioles with JF549 particle-tracks that met these criteria were then isolated and centred, and a custom Python script was used to calculate the distance between the centre of mass of the centriole (in the green channel) and the brightest pixel of the JF549 particle (in the orange channel) at successive timepoints.

SoRA super-resolution images were enhanced using Olympus Super-Resolution (OSR) algorithm with a filtering strength from low to standard, and subsequently deconvolved using a constrained iterative deconvolution algorithm with 5 iterations in CellSense software (Olympus).

### Protein structure prediction and visualisation

We initially screened for potential interactions between *Drosophila* AurA and Spd-2 using ColabFold v1.4.0 (Mirdita et al., 2022) in which AlphaFold2-multimer V2 was embedded to predict potential interaction interfaces between the full length proteins and/or various fragments of the proteins. Automatic model type was specified in run setup with number of recycles defined to 3 with no template information used. Using the iPTM score as an initial assessment of each predicted interaction, we identified the interaction between the AurA kinase domain (AurA155-421) and Spd-2291-310 (subsequently named ABD2) as the strongest hit (iPTM=0.769). Shortly afterwards, a potential interaction interface between HsAURKA and HsCEP192 was identified (Park et al., 2023). This was different to the potential interaction we initially identified, but in a screen using ColabFold v1.5.5 in which AlphaFold2-multimer V3 was embedded where we used the N-terminal half of Spd-2 (Spd-21-Spd-2650) with the AurA kinase domain (AurA155-421), an interaction interface similar to the one identified in humans was found with Spd-2229-250 (later named ABD1). Subsequent studies suggested that both ABD1 and ABD2 contributed to the recruitment of AurA to centrosomes, so we used AlphaFold2 ColabFold to predict the interaction interface between the AurA kinase domain and a combined Spd-2 peptide that contained both ABD1 and ABD2 (AurA155-421 and Spd-2229-310) (iPTM=0.59). PDB files for the human AURKA-Cep192 (PDB ID:8GUW) and AURKA-TPX2 (PDB ID: 5ODT) structures were downloaded from previously published structures on the PDB. UCSF Chimera X-1.7.1 (Pettersen et al., 2004) was used for structural analysis and figure generation.

### Immunoblotting

Embryos for immunoblotting were fixed and stored in methanol as described previously (Stevens et al., 2009). Afterward, the embryos were stored at 4°C at least overnight and rehydrated with 3× PBT (PBS + 0.1% Triton X-100) washes for 15 min each, before being subjected to electrophoresis and immunoblotting as described previously (Steinacker et al., 2022). The following primary antibodies were used: mouse anti-GFP (#11814460001, Roche) rabbit anti-TACC (1:1000) (Gergely et al., 2000b); rabbit anti-GAGA factor (1:500) (Raff et al., 1994); mouse anti-actin (A3853, Sigma-Aldrich); rabbit anti-GFP (A6455, Mol. Probes). HRP-conjugated donkey antirabbit (NA934V) and anti-mouse (NA931-1M (both from VWR International Ltd.) secondary antibodies were used at 1:3000-5000. Diluted SuperSignal ECL substrates (ThermoFisher Scientific) were used to develop chemiluminescence signal.

## Statistical analysis

The details of statistical tests, sample size, and definition of the centre and dispersion are provided in individual Figure legends.

## Code and Data availability

All raw image data will be deposited upon acceptance at The BioImage Archive (Hartley et al, 2022). The custom Python scripts used to process images can be downloaded from the Raff lab GitHub repository via https://github.com/RaffLab/.

---

**Figure S1.**
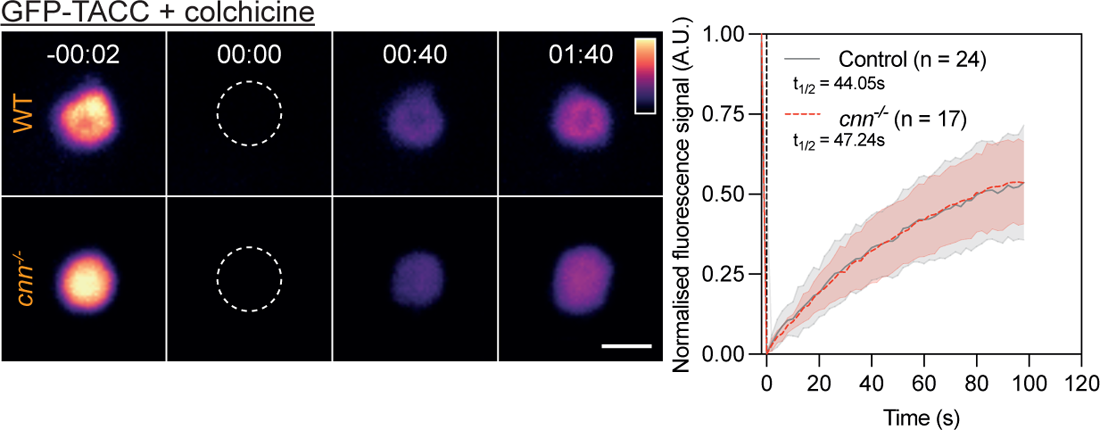
GFP-TACC recruitment to centrosomes does not appear to depend on Cnn. Images show, and graph quantifies, the centrosomal-fluorescence recovery (Mean±SD) after photobleaching of GFP-TACC in WT or *cnn^-/-^* embryos treated with colchicine. Time (mins:secs) is indicated; centrosomes were bleached at t=0:00. t_1/2_ was calculated from a fitted One-Phase Association Model (Graphad Prism). N=10-15 embryos, n=17-24 centrosomes. Scale bars = 2μm.

**Figure S2.**
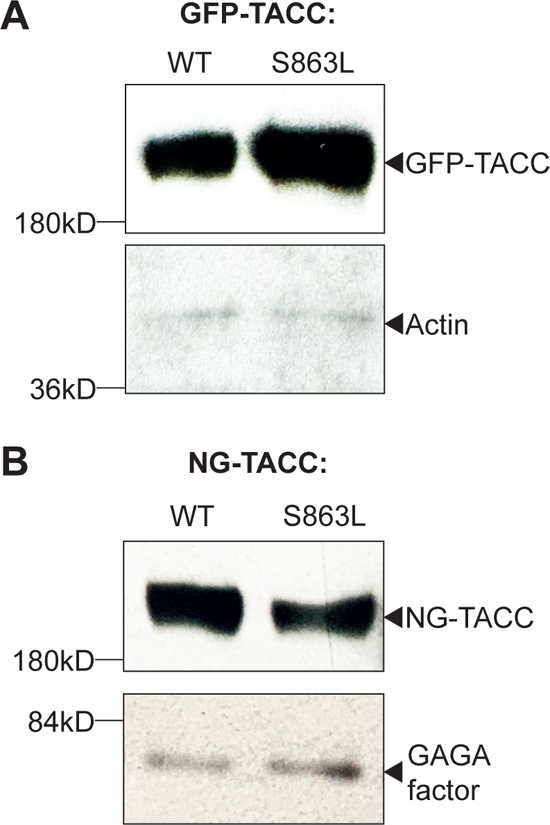
Expression levels of GFP- or NG-fusions to WT TACC or the TACC-S863L mutant. (A,B) 0-2hr old embryos expressing either WT GFP-TACC or GFP-TACC-S863L (A) (Barros et al., 2005) or WT NG-TACC or NG-TACC-S863L (generated in this study) (B) from the ubiquitin (Ubq) promoter were blotted with antibodies against either GFP (A) or TACC. Actin (A) or GAGA factor (B) (Raff et al., 1994) are shown as loading controls.

**Figure S3.**
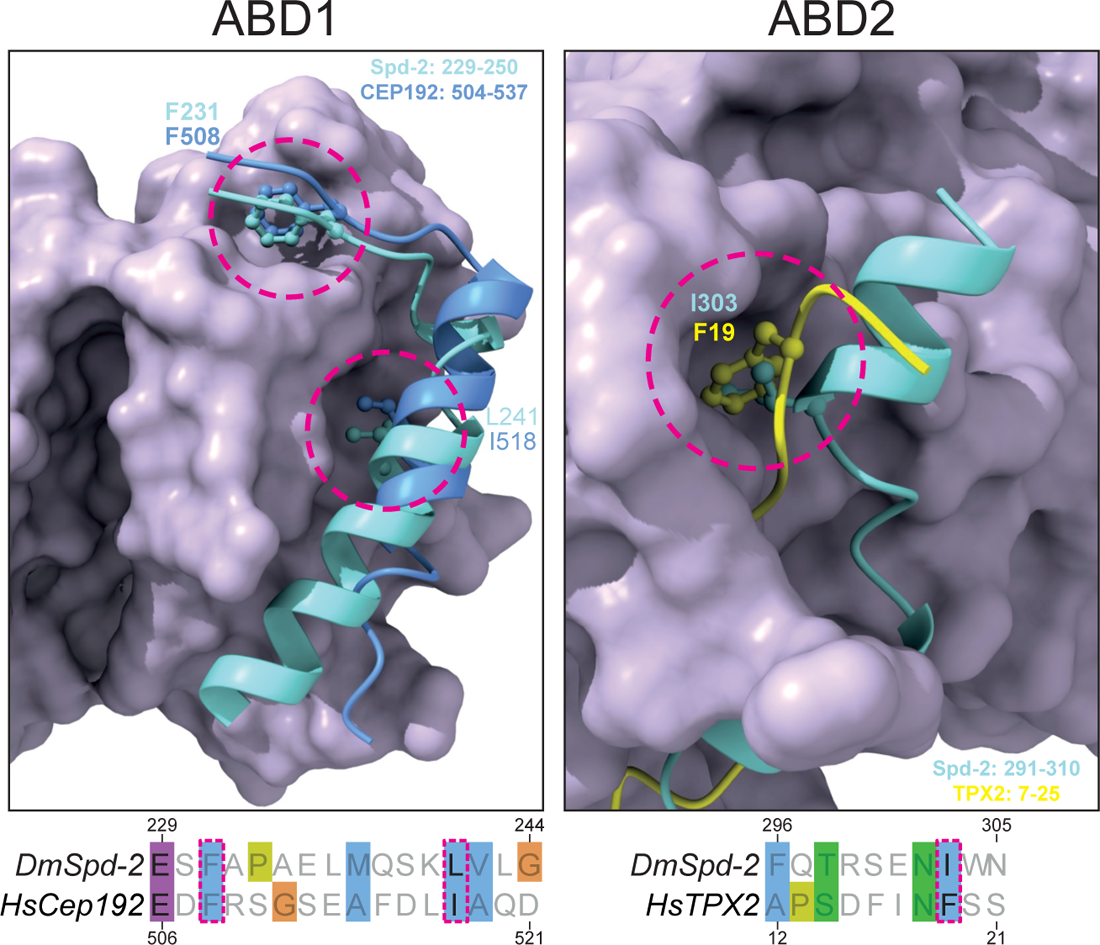
Analysis of the predicted structure of the Spd-2 AurA binding sites (ABD1 and ABD2). Images show a detailed view of the predicted interaction interface between the AurA kinase domain (magenta) and either ABD1 (cyan, left box) or ABD2 (*cyan*, right box), together with the crystal structure of the previously characterised region of either human CEP192 (*blue*, left box) or human TPX2 (*yellow*, right box) interacting with AURKA (but both shown here overlayed on the DmAurA structure). The sequences of the protein regions that interact with the AurA/AURKA kinase domain are aligned underneath the images (coloured using the Clustal Omega scheme). These sequences show limited conservation, but aspects of the binding of *Dm*Spd-2-ABD1 and *Hs*CEP192 to AurA/AURKA appear similar, and the position of two conserved hydrophobic amino acids that insert into the AurA/AURKA binding regions are highlighted with red-dotted lines. Interestingly, the substitution of the equivalent hydrophobic residues (F629 and I639) with charged Arg in Xenopus CEP192 strongly inhibits AurA binding (Joukov et al., 2014). Human TACC3 also interacts with this region of the AurA kinase domain (Burgess et al., 2018), with TACC3 F525 binding in a similar manner to F231 and F508 in the fly and human Spd-2/CEP192 proteins (not shown). Although DmSpd-2-ABD2 and HsTPX2_12-21_ clearly bind to a similar region on the AurA/AURKA surface, there is very limited similarity in how they do so—although a hydrophobic amino acid that inserts into a similar pocket on the AurA/AURKA surface is also highlighted in *red-dotted lines*.

**Figure S4.**
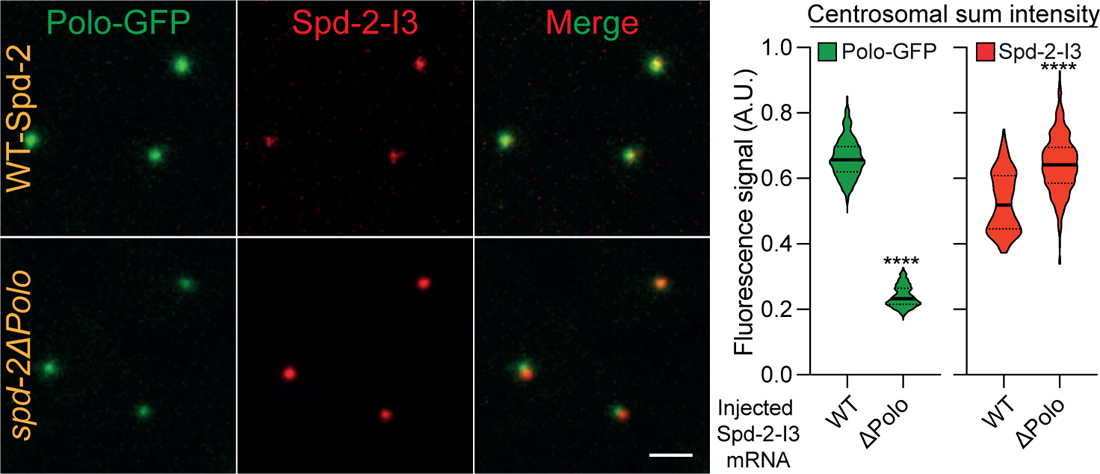
Images show, and violin plots quantify (Median±Quartile), centrosome fluorescence intensity in embryos expressing Polo-GFP and injected with mRNA encoding mScarlet-I3-fusions to either WT-Spd-2, or a mutant form of Spd-2 in which all 34 of the potential Polo-Box-Domain (PBD) binding motifs (S-S/T) were mutated to T-S/T, which prevents PBD-binding (Elia et al., 2003). The position of the 34 S-S/T motifs present in Spd-2 is indicated in Figure 5A (magenta lines). N= 10-11 embryos; n = 500-600 centrosomes for each group. Statistical significance was calculated using a Mann-Whitney (****: P<0.0001).

**Figure S5.**
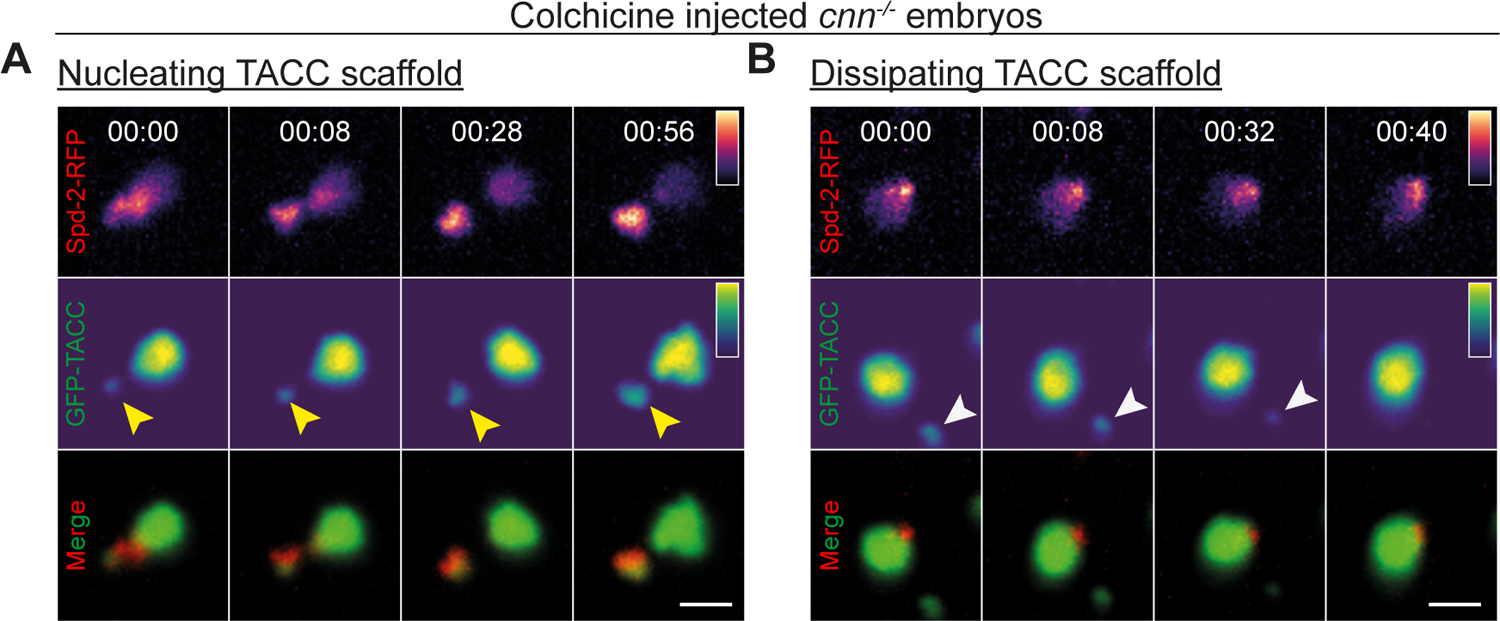
Centrioles appear to be required to generate the TACC scaffold. (A,B) Images from time-lapse movies show centrosomes in colchicine-injected cnn^-/-^ embryos co-expressing Spd-2-RFP and GFP-TACC. Time (mins:secs) is indicated. In (A) the centriole (yellow arrowhead) becomes separated from the main bulk of the TACC scaffold. New TACC scaffold starts to accumulate around the centriole. The centriole-less TACC scaffold remains stable over this short timescale, but will eventually start to dissipate (not shown). (B) This dissipation can be appreciated more easily at the centrosome shown here, where a smaller “flare” of GFP-TACC that lacks a centriole (*white arrowhead*) has become separated from the main TACC scaffold associated with the centriole. The flare lacks detectable Spd-2-RFP and it quickly dissipates. Scale bar = 2μm.

**Figure S6.**
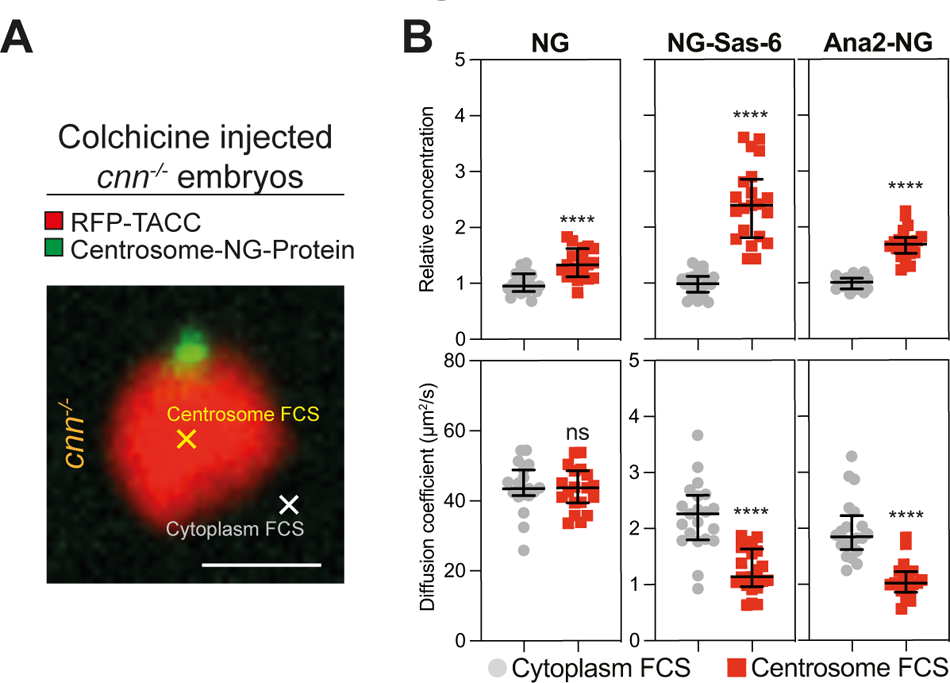
FCS comparison of protein behaviour at centrosomes in *cnn^-/-^* embryos. (A) Image shows a centrosome in an embryo expressing RFP-TACC and Ana2-NG and injected with colchicine. This illustrates the typical areas that were analysed by FCS in the outer regions of the centrosome (*yellow cross*) or the nearby cytoplasm (*white cross*). In the absence of Cnn scaffold, the PCM appears to be structurally weakened and the centriole cannot maintain its position at the centre the PCM (Lucas and Raff, 2007). Scale bar = 2μm. (B) Scatter plots show the FCS-measured concentration (top plots) or diffusion rate (bottom plots) (Median±Quartiles) of NG or various NG-fusions in the cytoplasm (*grey circles*) or in the outer regions of the centrosome (*red squares*) in WT embryos injected with colchicine. N=20-25 embryos. Statistical significance was calculated using Mann-Whitney’s test (****: P<0.0001, ns: not significant).

